# OSBP-mediated cholesterol transfer determines epithelial polarity and associated cargo secretion

**DOI:** 10.1101/2021.12.09.471984

**Authors:** Dávid Kovács, Anne-Sophie Gay, Lucile Fleuriot, Delphine Debayle, Ana Rita Dias Araújo, Amanda Patel, Bruno Mesmin, Frédéric Luton, Bruno Antonny

**Affiliations:** CNRS, Université Côte d’Azur, Institut de Pharmacologie Moléculaire et Cellulaire, Valbonne, France

**Keywords:** Membrane contact sites, Lipid transfer proteins, cargo secretion, OSBP, cell polarity, EMT

## Abstract

Golgi lipid environment regulates sorting and cargo secretion. However, the mechanisms that spatiotemporally control the lipid composition of the secretory membranes to drive cargo trafficking are poorly understood. Lipid transfer proteins regulate the concentration of specific lipids at membrane contact sites. We hypothesised that by catalysing cholesterol/PI(4)P exchange at ER-*trans*-Golgi membrane contact sites the lipid transfer protein oxysterol binding protein (OSBP) affects the secretion of a subset of cargoes. Here, we report that OSBP is a major epithelial protein as its inhibition leads to complete loss of apico-basal polarity. By mapping the OSBP proximity proteome with the biotin ligase TurboID, we found that OSBP controls the secretion of multiple membrane associated proteins, including key polarity determinants such as E-cadherin. Mechanistically, we established that OSBP contributes to E-cadherin secretion by supplying cholesterol to post-Golgi membranes. Importantly, when cells downregulate cell-cell junctions upon epithelial-to-mesenchymal transition, they re-wire their lipid homeostasis and downregulate OSBP as well, thus altering the trafficking of the OSBP-dependent secretory cargoes.

## Introduction

In eukaryotic cells, the Golgi apparatus is responsible for the sorting and the secretion of about 30% of the neosynthesized proteins (Boncompain & Weigel, 2018). Although Golgi sorting signals and their recognizing machineries are well-characterised, the molecular mechanisms of clustered cargo secretion are still largely unknown (Ramazanov et al., 2021; Boncompain & Weigel, 2018). It has already been long recognised that lipid environment affects protein secretion (Keller & Simons, 1997, 1998). Indeed, lipid composition defines the biophysical properties of the membranes, which might facilitate the formation of different Golgi sub-domains and the recruitment of various proteins leading to the assembly of cargo-specific trafficking machineries (Blume & Hausser, 2019; Klemm et al., 2009). However, our understanding of the link between lipid composition and membrane trafficking has remained fragmental due to the difficulty in tuning specifically and locally the lipid composition of organelles.

Recent advances in the characterization of membrane contact sites (MCS) offer new strategies to study the impact of lipid composition on membrane traffic steps. MCS, which are regions of close apposition between organelles, are enriched in lipid transfer proteins (Wu et al., 2018). Oxysterol binding protein (OSBP) – a member of the OSBP-related protein family (ORPs) – is recruited to the ER-Golgi MCS to transfer cholesterol from the ER towards the *trans*-Golgi network (TGN), thereby controlling the lipid composition of the *trans*-Golgi secretory membranes (Mesmin et al., 2013, 2019). This transfer activity requires the biochemical energy of the lipid phosphatidylinositol-4-phosphate (PI(4)P), which has an unequally distributed concentration between ER and Golgi membranes (Antonny et al., 2018; Mesmin et al., 2013, 2017). While PI(4)P concentration is high in the *trans*-Golgi due to the activity of Golgi-resident PI4-kinases, it is degraded in the ER by the phosphatase Sac-1. When OSBP transfers a cholesterol molecule to the Golgi, it extracts a PI(4)P from the Golgi membrane and transfers it towards the ER. Thus, cholesterol is exchanged for PI(4)P during OSBP-mediated lipid transfer (Mesmin et al., 2013, 2017). Because cholesterol and PI(4)P are recognised as major lipids controlling trafficking and secretion (Blume & Hausser, 2019; Dukhovny et al., 2009; Sugiki et al., 2012), OSBP might affect the trafficking route of cargoes that depend on the local amount of these lipids.

OSBP deletion in mice is lethal at early stages of embryogenesis demonstrating its essential function (Brown and Goldstein, personal communication). Indeed, OSBP is highly expressed in most human tissues, and the fact that virtually all of our cells are able to synthesise and distribute cholesterol suggests that OSBP-catalysed lipid transfer is universal in the body. However, OSBP targeting drugs such as Schweinfurthin G (SWG) trigger cytotoxicity in a cell line-specific manner with cell line sensitivity to SWG being inversely proportional to cellular OSBP levels (Burgett et al., 2011; Péresse et al., 2020; Rajapakse et al., 2018). These variations in OSBP expression levels and sensitivity to OSBP-specific drugs suggest that cells with distinct phenotypes might require different amounts and/or activity of OSBP. However, it is unclear which cell types need elevated OSBP and for which functions.

Epithelial cells line the surfaces of our body and are differentiated to form tightly-connected cellular sheets that act as biological barriers. To achieve this function, epithelial cells are polarized on their apico-basal axis and express high levels of cell junction components such as cadherins (Du et al., 2010; Moreno-Bueno et al., 2006; Shigetomi & Ikenouchi, 2019). On the other hand, mesenchymal cells are not polarised on an apico-basal axis and due to differences in cadherin expression, they are not able to form stable cell-cell junctions, which allows them to gain migratory potentials and leave their original environment (McFaline-Figueroa et al., 2019). The process by which an epithelial cell gradually loses its phenotype and acquires mesenchymal features is referred to as epithelial-to-mesenchymal transition (EMT) (Carmona et al., 2014; Karimi Roshan et al., 2019; Loh et al., 2019). Although the EMT program is part of many physiological processes such as embryonic development or wound healing, it is often related to cancerous malignancies since EMT is frequently observed in cancer cells and can facilitate metastatic dissemination (Jia et al., 2016; Morandi et al., 2017; Ocaña et al., 2012). Therefore, identifying new aspects of the EMT program is crucial for future therapeutic developments.

Here, we demonstrate that OSBP at ER-Golgi contact sites regulates the secretion of a set of cargo proteins in epithelial cells, such as cell junction proteins including E-cadherin, thereby contributing to the maintenance of epithelial polarity. Moreover, OSBP-dependent cadherin trafficking relies on the cholesterol-transfer activity of OSBP. Importantly, when cells downregulate cell-cell junctions during EMT, OSBP-mediated lipid transfer in the ER-TGN contact sites is also reduced, demonstrating the plasticity of the OSBP-regulated secretory pathway upon this phenotypic switch.

## Results

### 1. Epithelial phenotype is associated with high OSBP expression

Epithelial cells are typically polarized along an apico-basal axis and need to organize their secretory routes to direct cargo molecules according to this geometry. To test whether members of the ORP family contribute to this function, we first performed a bioinformatics screen to examine whether the expression of OSBP and the other members of the ORP family varies across cell lines with different epithelial characteristics (Rajapakse et al., 2018). To this aim, we extracted the gene expression data of the cell lines included in the Cancer Cell Line Encyclopedia (Barretina et al., 2012) focusing on all ORP genes as well as a panel of epithelial and mesenchymal marker genes (**Fig 1A**). To define a cell-line specific epithelial-mesenchymal signature, we calculated a single epithelial-mesenchymal index (EMI) for each cell line using the marker gene expression data (**Fig 1B**) and we correlated these values with the expression levels of the various ORPs (**Fig EV1A, Fig 1C**). This analysis indicates that OSBPL2 and OSBP expression levels correlate positively with EMI, while OSBPL8 and OSBPL6 expressions show a negative correlation. Thus, OSBPL2 and OSBP are highly expressed in cell lines with epithelial characteristics, while OSBPL8 and OSBPL6 are rather mesenchymal cell-specific genes.

**Figure 1.**
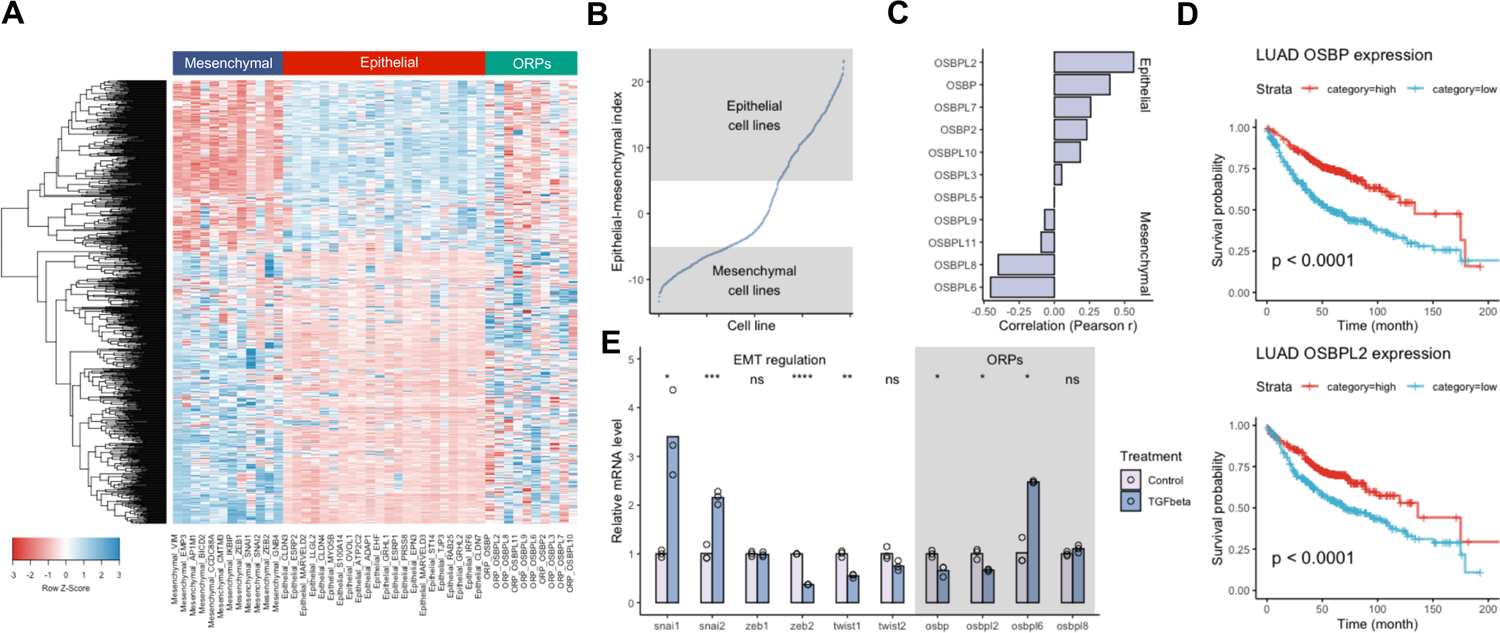
OSBP is highly expressed in epithelial cells. A. Expression heatmap of epithelial, mesenchymal and ORP genes across cell lines of the Cancer Cell Line Encyclopedia collection.
B. Calculated epithelial-mesenchymal indexes corresponding to each cell line.
C. Correlation between epithelial-mesenchymal index and ORP expression reveals epithelial and mesenchymal-specific ORPs.
D. Low OSBP and OSBPL2 expression are associated to a poor clinical outcome in lung adenocarcinoma patients. P-values were calculated by Gehan-Breslow-Wilcoxon test. LUAD: Lung adenocarcinoma.
E. TGFbeta-induced expression changes in master EMT-transcription factors and selected ORP genes measured by RT-qPCR. Means are calculated from three independent experiments. *p<0.05; **p<0.01; ****p<0.001; ns=non-significant, Welch’s t test.

To test whether this observation can be extended to disease conditions as well, we repeated the analysis using RNAseq data obtained from TCGA lung adenocarcinoma specimens (**Fig EV1B-E**). In agreement with the cell-line collection data, we found that OSBP expression positively correlates with the epithelial feature of the given cancer tissue specimens. EMT can lead to metastatic dissemination, therefore tumours with mesenchymal characteristics are often coupled to a shorter patient survival. By analysing lung adenocarcinoma survival data, we found that high OSBP and OSBPL2 expressions are associated to significantly longer patient survivals further suggesting that downregulation of these genes is a signature of tumour malignancy (**Fig 1D**).

To experimentally test the re-distribution of ORP gene expression pattern upon EMT, we triggered EMT in A549 lung adenocarcinoma cells. Although this cell line shows both epithelial and mesenchymal characteristics, its mesenchymal phenotype can be further enhanced by TGFbeta treatment (Hajek et al., 2012; Hao et al., 2019; Hua et al., 2019). We assessed the relative expression of EMT transcription factors and the top-changed ORP mRNA levels by RT-qPCR. While snai1 and snai2-inductions dominated the TGFbeta-triggered EMT in A549 cells, we observed a concomitant decrease in OSBP and OSBPL2. A significant increase in OSBPL6 messengers was measured, validating our previous findings, whereas no change in OSBPL8 expression was observed (**Fig 1E**).

To further investigate the epithelial expression of OSBP and OSBPL2, we meta-analysed histology samples of the Human Protein Atlas (Uhlen et al., 2015). As revealed by specific antibodies, OSBP and OSBPL2 proteins exclusively decorate the epithelial cell layers of various tissue samples such as nasopharynx and urinary bladder, while the skeletal muscle, which has a mesenchymal origin, lacks OSBP and expresses high levels of OSBPL6 (**Fig EV1F**).

We concluded that among all ORP genes, OSBP and OSBPL2 are prominent epithelial proteins, showing high expression in cell lines with epithelial features and down-regulation in response to TGFbeta. In the following experiments, we focused on OSBP since its lipid transfer activity is well-characterized in cells and can be targeted by specific drugs, allowing us to precisely dissect its potential involvement in epithelial secretion regulation.

### 2. OSBP inhibition disrupts epithelial polarity

To test whether silencing of OSBP expression or its targeting with pharmacological inhibitors affect cell polarity establishment and maintenance, we used MDCK cells, which are able to form cysts with polarized cells when cultured in extracellular matrix (Mangoo-Karim et al., 1989; Qin et al., 2010). MDCK cells were nucleofected with siRNA targeting OSBP, then embedded into Matrigel 24 h later and left to establish cysts with polarized cells for 3 days. Efficient OSBP silencing was confirmed in cells lysed 4-days after nucleofection (**Fig 2B**). The developed cysts showed a typical morphology where the apical surface of the cells is oriented toward a single central open lumen. The tight junction marker ZO-1 decorated the intersection of the apical and lateral plasma membrane surfaces, E-cadherin was found in the basolateral domain, and Gp135 localised exclusively on the apical surface of the cells (**Fig 2A**). While only ∼35% of the cysts showed irregular morphology in the control siRNA-nucleofected cells, ∼50% of the OSBP silenced cyst population showed luminal morphology defects (**Fig 2A-C**). Typically, these cysts developed multiple lumens and displayed apical multipolarity where one cell formed several apical surfaces.

**Figure 2.**
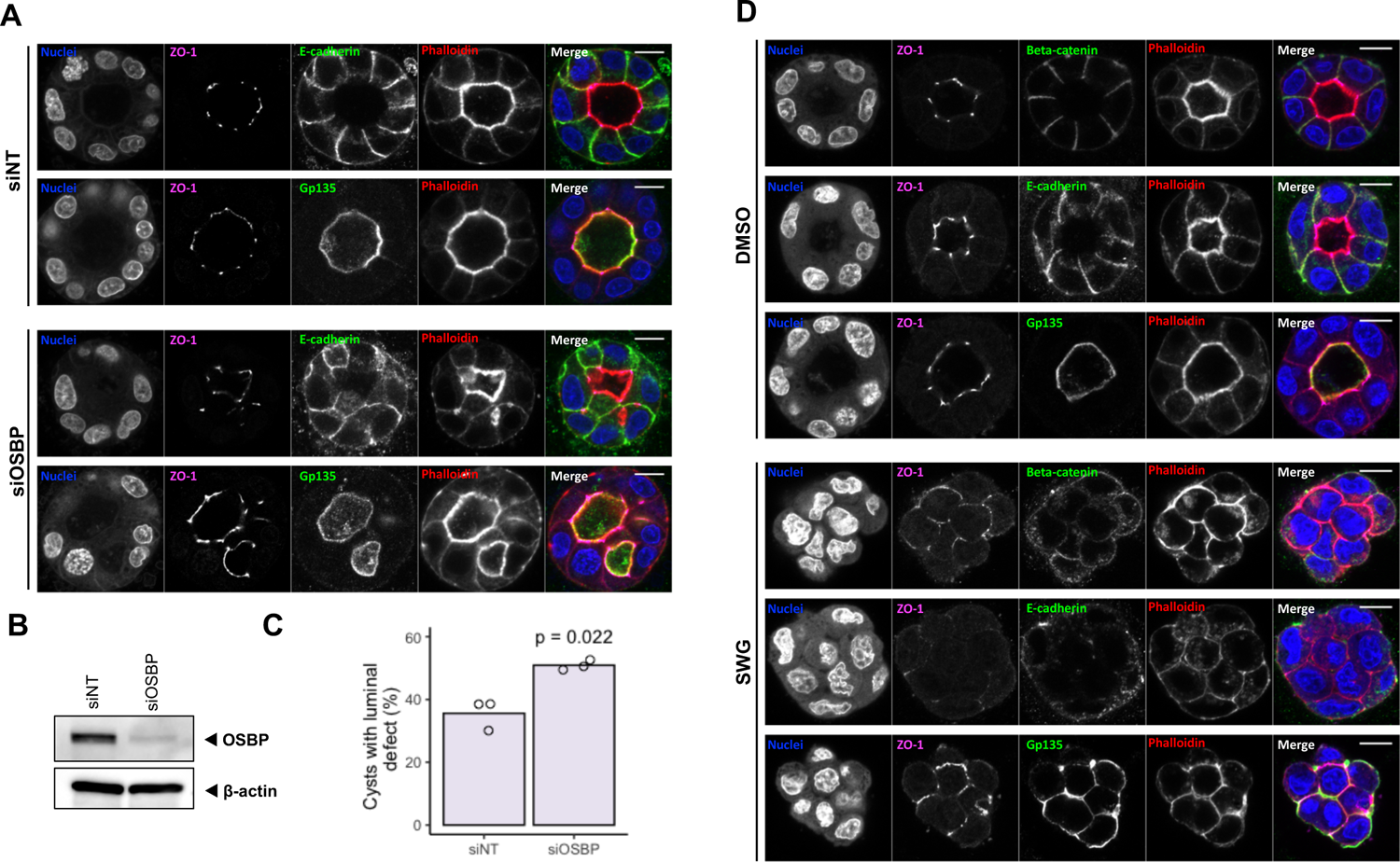
OSBP regulates epithelial polarity. A. OSBP silencing affects MDCK cyst morphology. Representative cysts stained with apical/basolateral markers are shown. Bar=10 #m
B. Efficacy of RNAi-based OSBP silencing was tested 4 days post-nucleofection in MDCK cells.
C. OSBP silencing increases the frequency of MDCK cysts with morphology defects. Means were calculated from three independent experiments. Each experiment was done in triplicates/condition and statistics was obtained by counting at least 70 cyst/triplicate. P-value was calculated by unpaired t test.
D. Overnight treatment of preformed MDCK cysts with 0.5 μM SWG disrupts polarized morphology. Representative cysts stained for apical/basolateral markers are shown.

To test the effect of OSBP inhibition on polarized cells, we next used SWG, a drug belonging to the ORPphilin family and that it is very specific for OSBP (Burgett et al., 2011; Péresse et al., 2020). Strikingly, overnight treatment of polarized MDCK cysts with the OSBP-targeting drug SWG leads to a complete loss of cell polarity, as the lumens disappeared and the cysts acquired a grape-like morphology consisting of round-shaped cells showing no asymmetrical marker distribution (**Fig 2D**). Although no apical and basolateral membrane domains could be distinguished in SWG-treated cysts, ZO-1 and Beta-catenin-positive membrane segments showed no overlap, suggesting that these domains were able to partially preserve their identity upon OSBP blockage. Interestingly, in SWG-treated cysts, a reduced Beta-catenin and E-cadherin signal was detected at cell-cell junctions. Additionally, Gp135 was detected in plasma membrane domains facing towards the extracellular matrix as well (**Fig 2D**). Altogether, the gene silencing and pharmacological approaches indicated that a functional OSBP is essential for the establishment as well as for the maintenance of the epithelial polarity.

### 3. OSBP proximity landscape reveals epithelial polarity determinants close to ER-TGN contact sites

OSBP might contribute to epithelial polarity maintenance by regulating post-Golgi transport pathways through lipid transfer between ER and TGN membranes. To test this hypothesis, we decided to identify secretory cargoes that are localised in the physical proximity of OSBP. We chose a proximity biotinylation strategy in which the promiscuous biotin ligase TurboID was fused to the C-terminus of OSBP (**Fig EV 2A**). When expressed in cells, TurboID biotinylates proteins that are within a 20 nm radius of the fusion protein (Branon et al., 2018). The biotinylated proximity partners are then purified and identified by mass spectrometry.

In MDCK cells stably expressing OSBP-TurboID, the fusion bait was detected both in the cytoplasm as well as bound to TGN-marker bGalT1 positive Golgi structures (**Fig 3A**). Accordingly, biotinylated proteins revealed by fluorescently-labelled streptavidin were detected in the cytoplasm as well as in the Golgi of OSBP-TurboID expressing MDCK cells. Next, we treated the cells with OSBP inhibitors SWG and OSW-1 for 60 min prior to biotinylation to increase the partitioning of OSBP to ER-TGN contact sites at the expense of the cytosolic pool (Burgett et al., 2011; Mesmin et al., 2017; Péresse et al., 2020). Under these conditions, we observed a drop in the cytoplasmic streptavidin signal and a concomitant increase in the TGN-localised OSBP pool. Thus, the OSBP-TurboID bait protein behaved as its endogenous counterpart (**Fig 3A-B**). In addition, ORPphilin treatments promoted the apparition of OSBP-TurboID on round-shaped cytoplasmic structures, confirming that a pool of OSBP can be recruited to endo-lysosomal surfaces as well (Lim et al., 2019).

**Figure 3.**
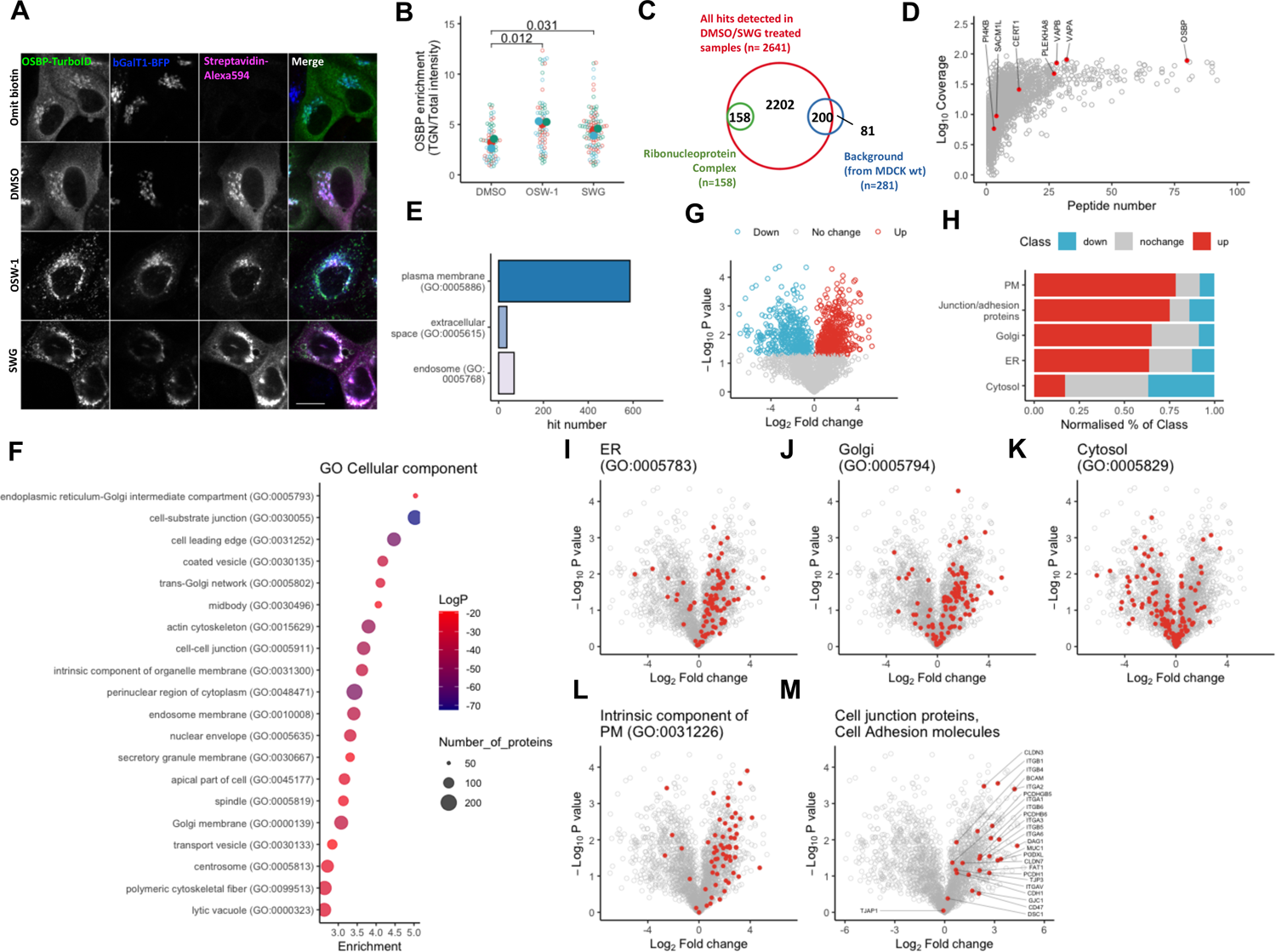
OSBP proximity landscape highlights secretory cargoes depending on ER-TGN contact sites. A. Confocal microscopy shows the OSBP-TurboID-derived biotinylation signal in MDCK cells. Cells stably expressed bGalT1-BFP to label TGN. OSW-1 was used in 50 nM and SWG was applied in 0.5 μM concentration for 1 h. Bar=15 #m.
B. Superplot showing TGN-enrichment of OSBP-TurboID upon OSW-1 and SWG treatments. Means obtained from three independent experiments were compared by Welch’s t test.
C. Venn diagram showing the filtering of the identified hits detected by mass spectrometry.
D. Numerous well-characterized OSBP-partners were identified in the OSBP proximity proteome. See complete list in **Source Data 1**.
E. Classification of identified hits by cellular components shows that a large portion of the OSBP-proximity proteins are localised to the plasma membrane.
F. GO cellular component enrichment analyses shows significant enrichment in multiple GO classes.
G. Classification of identified hits according to their response to 1 h 0.5 μM SWG treatment.
H. Enrichment of hits belonging to specific GO classes shows different profiles to SWG treatment.
I. (I-M) Mapping hits belonging to indicated GO classes shows different distributions across the proximity interactome.

For proteomic analysis, we exposed MDCK cells stably expressing OSBP-TurboID to either DMSO or SWG for 60 min followed by biotin addition during the last 10 min of treatment (**Fig EV2B**). We verified successful protein biotinylation and purification by streptavidin-HRP blotting and silver staining of the elution fractions, respectively (**Fig EV2C-D**). Following on-bead trypsinization, peptides were subjected to mass spectrometry and proteins were identified by subsequent proteomics analyses. Overall, we identified 2641 proteins by this strategy, among which 200 were considered as background since these hits were also identified on beads subjected to lysates from biotin-treated MDCK cells not expressing OSBP-TurboID (**Fig EV2B**). As the nascent TurboID polypeptide can already catalyse protein biotinylation, proteins classified by the Gene ontology (GO) group as “Ribonucleoprotein Complex” were also considered as false positive hits and excluded from further analysis (**Fig 3C**).

In the DMSO-treated cells, we identified many proteins already known to interact or co-localise with OSBP at MCS. These include the VAP isoforms (VAPA and VAPB), the PI(4)P generating kinase PI4KIIIbeta (PI4KB) and the ER-resident phosphatase SAC1 (SACM1L). Additionally, we detected the ceramide transporter CERT1 and the glucosylceramide-transfer protein FAPP2 (PLEKHA8), confirming their collaboration with OSBP at the ER-TGN contact sites (**Fig 3D**).

Next, we wanted to identify potential secretory cargoes among the identified hits. Since Golgi secretion directs cargoes to the extracellular space, to the endo-lysosomal system or toward the plasma membrane, we classified the hits according to their final cellular localization. We identified 37 proteins listed in the GO group “extracellular space”, 69 categorized as “endosome”, and 586 classified as “plasma membrane” (**Fig 3E**), suggesting that the majority of the OSBP-proximity cargoes are proteins directed toward the plasma membrane. Furthermore, we found a large overlap between previously published MDCK surface proteome (Caceres et al., 2019) and the OSBP proximity proteome, suggesting that a considerable portion of the surface proteins are getting close to OSBP upon their secretion (**Fig EV2E**). However, when we correlated the abundance of identified proximity partners in the DMSO-treated cells with the apico-basal localization of the MDCK surface proteins, we established no correlation, indicating that OSBP has no preference toward apical or basolateral cargoes (**Fig EV2F**).

To further characterize the proteins of the OSBP proximity landscape, we performed a statistical analysis using Metascape (Zhou et al., 2019), which allowed us to assess significant enrichments in cellular components, molecular function and biological processes-related GO terms. As expected, statistically significant enrichments were found in multiple GO groups encompassing ER and Golgi-resident proteins. Importantly, other GO terms such as “cell-substrate junction” and “cell-cell junctions” showed significant enrichments as well (**Fig 3F**). Strikingly, “cadherin binding” and “establishment or maintenance of cell polarity” GO groups were found amongst the top significantly enriched groups upon classification by molecular function and biological processes (**Fig EV2G-H, Source Data 1**). Besides the numerous adherent junction components (e.g. cadherins and catenins), we identified tight junction proteins (e.g. TJP1 and 2, Occludin, MarvelD2 and Claudin), integrins, CD44, EpCam and the apical marker proteins PODXL and MUC1, all of which are regulators or determinants of epithelial polarity, suggesting that OSBP contributes to epithelial polarity by regulating these factors.

Because SWG-treatment stabilizes OSBP at contact sites, cargoes close to ER-TGN contact sites should show an increased abundance among the hits when cells are treated with this drug. Thus, we divided the hits into three groups according to their fold change upon SWG treatment: hits that showed no significant change, hits with increased and hits with decreased abundance across the OSBP proximity proteome upon treatment (**Fig 3G**). Then, we mapped specific classes of hits to the volcano plot to show their SWG-dependent distribution in the OSBP proximity landscape. Hits classified as ER and Golgi-resident factors were enriched with the highest extent among proteins that showed an increased abundance around OSBP after SWG treatment (**Fig 3H-J**). On the contrary, most cytosolic hits showed a decreased abundance in the vicinity of OSBP when cells were treated with SWG (**Fig 3H, K**). Next, we mapped proteins that are classified as “intrinsic components of the plasma membrane” and, thereby, potential secretory clients of OSBP. The majority of these proteins showed an increased abundance upon SWG-treatment in the OSBP proximity proteome (**Fig 3H, L**). Since the enrichment analyses indicated the presence of many cell junction components among the identified partners, we wanted to see whether their abundance in the OSBP proximity proteome changes upon SWG-treatment as well. For this, we selected those hits that are classified as cell “junction proteins” or “cell adhesion molecules” and we found that the majority of these proteins enriched around OSBP upon SWG-treatment (**Fig 3H, M**). Thus, a high number of cell surface proteins and cell junction molecules become more abundant in the proximity of OSBP when it is forced to accumulate at ER-Golgi contact sites by SWG-treatment, indicating that these proteins become close to OSBP when they are sorted in the Golgi membranes along their secretion.

### 4. OSBP controls E-cadherin trafficking and cell surface expression

Together with numerous cell junction components (**Source Data 1**), E-cadherin was identified in the OSBP proximity, suggesting that OSBP contributes to its trafficking. E-cadherin expression on the cell surface is one of the major hallmarks of epithelial organization, and its loss is frequently observed in carcinoma demonstrating that it functions as a potent tumour suppressor (Dongre & Weinberg, 2019; Thiery, 2002). Thus, we decided to use E-cadherin to validate the functional interaction between OSBP and one of its identified proximity partners. Immunoblot analysis of whole cell lysates and pull-down of biotinylated proteins obtained from OSBP-TurboID-expressing MDCK cells showed that only a minor portion of the total E-cadherin pool got biotinylated over the 10 min-biotin treatment, suggesting transient interaction between OSBP and E-cadherin (**Fig EV3A**). Remarkably, incubating the cells for 2 h in biotin-free medium after a 10 min biotin exposure led to the appearance of the biotinylation signal at the cell-cell junctions, whereas no cell-boundary-specific signal was observed when the cells were fixed directly after the 10 min-biotin treatment (**Fig 4A**). Hence, we hypothesised that neosynthesized E-cadherin gets into the proximity of OSBP along its sorting in the *trans*-Golgi network.

**Figure 4.**
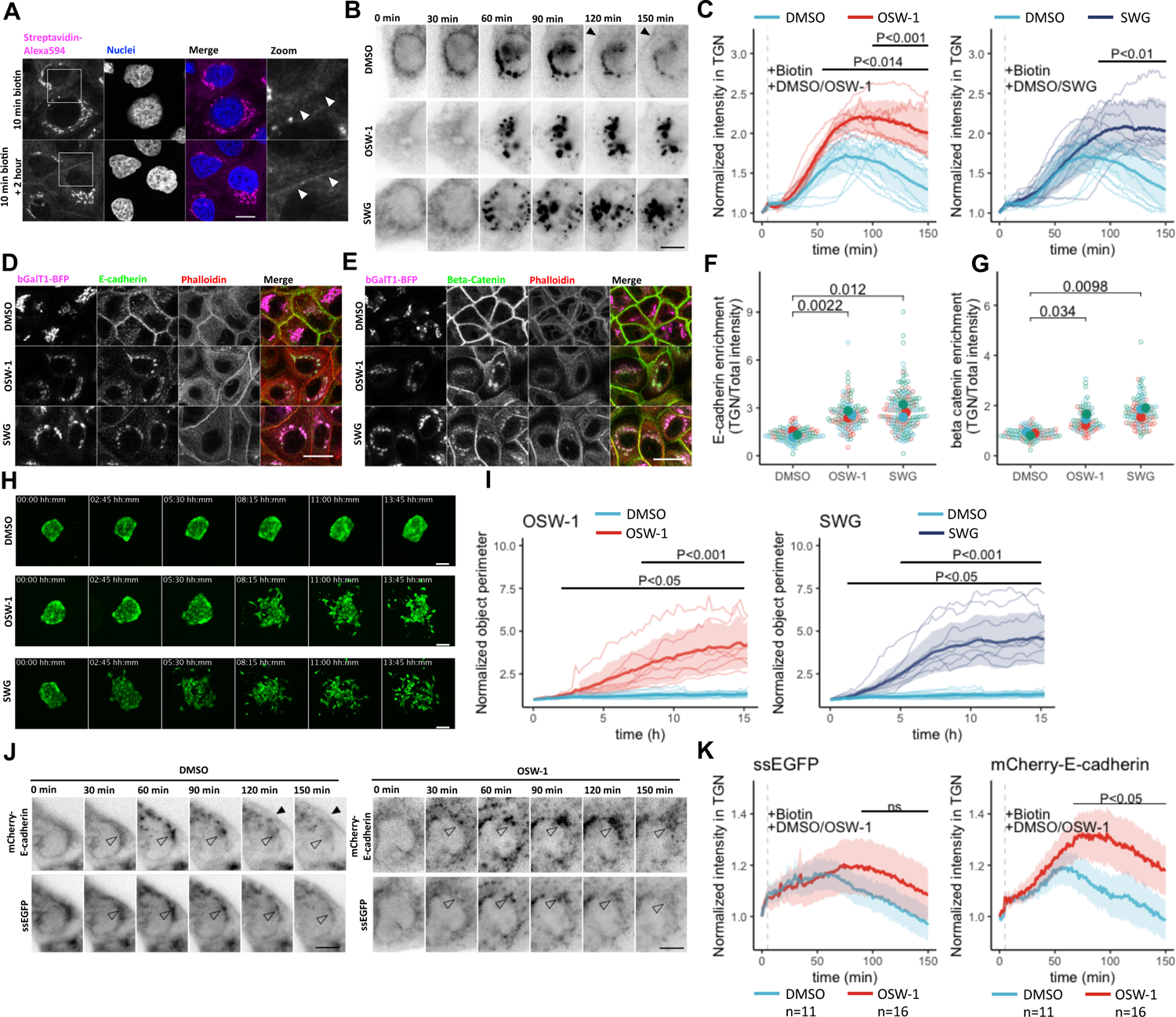
OSBP controls E-cadherin trafficking. A. Confocal imaging shows that incubating OSBP-TurboID expressing MDCK cells for 2 h after biotin treatment results in the enrichment of biotinylation signal at cell-cell junctions (arrowheads). Bar=12 #m
B. Live epifluorescent imaging shows the arrest of E-cadherin secretion at the Golgi membranes upon ORPphilin treatments (0.5 μM SWG and 20 nM OSW-1). Black arrowheads points to cell-surface localised E-cadherin. Bar=10 #m
C. Kinetics of the E-cadherin RUSH experiments. One representative experiment is shown. Each line corresponds to one cell and bold lines with the shaded area indicate mean±95% confidence interval. P-values were calculated by unpaired t test.
D. Immunolabelling of endogenous E-cadherin observed by confocal microscopy. Bar=20 #m.
E. Immunolabelling of endogenous beta-catenin observed by confocal microscopy. SWG was used in 0.5 μM and OSW-1 in 2 nM upon experiments shown on both D and E. Bar=20 #m.
F. Superplot showing E-cadherin enrichment in the TGN upon ORPphilin treatment. Means obtained from three independent experiments were compared by Welch’s t test.
G. Superplot showing beta-catenin enrichment in the TGN upon ORPphilin treatment. Means obtained from three independent experiments were compared by Welch’s t test.
H. Representative colonies showing the effect of ORPhilin treatments (0.5 μM SWG and 2 nM OSW-1)on cell scattering. Bar=60 #m
I. Cell scattering was quantified by measuring colony perimeters. Each line corresponds to one colony and bold lines with the shaded area indicate mean±95% confidence interval. P-values were calculated by unpaired t test.
J. Co-expressed mCherry-E-cadherin and ssEGFP rush cargoes in MDCK cells show different sensitivities toward 20 nM OSW-1 treatment. Empty arrowheads indicate Golgi membranes while black arrowheads point to surface-localised E-cadherin. Bar=10 #m
K. Quantification of the co-expression experiment. One representative experiment is shown. Each line indicates mean±95% confidence interval. DMSO n=11, OSW-1 n=16. P-values were calculated by unpaired t test.

To directly assess whether E-cadherin trafficking depends on OSBP, we applied the retention using selective hooks (RUSH)-system using EGFP-tagged E-cadherin as a model cargo (Boncompain et al., 2012). This method consists of trapping a chosen cargo by streptavidin-dependent interaction with ectopically co-expressed ER-resident KDEL, and in promoting a wave of cargo secretion by adding biotin to compete out the E-cadherin/KDEL interaction. With this system, we observed that E-cadherin was released from the ER and transported to the Golgi within 60 min after biotin addition (**Fig 4B, C**). Thereafter, E-cadherin left the Golgi and appeared at the cell-cell junctions in ca. 120 min (**Fig 4B** arrowheads, **Movie EV1**). When biotin was co-administrated with OSBP inhibitor OSW-1 or SWG, the kinetics of ER to Golgi transport of E-cadherin was comparable to that of the control. In contrast, the exit of E-cadherin from the Golgi apparatus was significantly delayed upon both OSW-1 and SWG treatments, resulting in higher E-cadherin signal at the Golgi (**Fig 4B, C, Movie EV1**). Thus, OSBP function is essential for complete E-cadherin secretion.

To test whether OSBP blockage affects the secretion of endogenous E-cadherin as well, we treated TGN-marker bGalT1-BFP-expressing MDCK cells with OSW-1 or SWG and fixed the cells 4 h after treatment to immunolabel endogenous E-cadherin (**Fig 4D**). As expected, E-cadherin in the control samples decorated mostly the cell-cell junctions, whereas it disappeared from the cell-cell junctions in the treated samples and accumulated in the *trans*-Golgi network (**Fig 4D, F**). Importantly, we observed the same trend when we investigated beta-catenin – another cargo identified in the proximity biotinylation screen – highlighting that other, non-transmembrane components of adherent junctions follow the same OSBP-dependent trafficking route (**Fig 4E, G**).

To further test the role of OSBP on adherent junction regulation, we grew EGFP-expressing MDCK cells to form colonies and performed overnight time-lapse imaging after the addition of the OSBP-inhibiting drugs OSW-1 and SWG (**Fig 4H-I, Movie EV2**). In the DMSO-treated control, we observed only basal expansion of the colonies due to cell proliferation. However, after 8 hours exposure to ORPphilins, the colonies started to scatter, indicating that OSBP-inhibition has a striking impact on cell-cell junction dynamics.

To test whether secretion of other cargo types also show dependency on OSBP, we applied the RUSH system in cells co-expressing mCherry-tagged E-cadherin cargo with small soluble, luminally retained EGFP (ssEGFP). Following biotin-induced release, both model cargoes entered the Golgi network and followed similar secretory kinetics (**Fig 4J-K**). Co-administrating OSW-1 with biotin, however, triggered only a minor shift in soluble EGFP-secretion, while this treatment affected mCherry-E-cadherin secretion with a higher degree (**Fig 4K, Movie EV3**). In line with previous observations on other model cargoes (Boncompain et al., 2012), confocal microscopy showed that E-cadherin and ssEGFP diverge in their trafficking as non-overlapping EGFP and mCherry-E-cadherin regions were observed 60 and 120 min following biotin administration (**Fig EV3B**). This experiment shows that OSBP inhibition selectively affects the trafficking of distinct cargoes.

### 5. OSBP-driven cholesterol is necessary for epithelial polarity and cargo secretion

Because OSBP loads TGN membranes with cholesterol, we hypothesised that the previously observed effects of OSBP inhibition on cargo trafficking is due to drop of cholesterol level along the secretory membranes. To test this, we first investigated the effect of OSBP silencing on the distribution of cholesterol in 3D polarized MDCK cells. As expected, the probe D4H-GFP, which recognizes accessible cholesterol in membranes (Das et al., 2014; Maekawa, 2017; Takahashi et al., 2021), decorated cell membranes of the polarized cells and an enrichment of the fluorescent signal was detected at the basolateral domain of the plasma membrane of the control siRNA-transfected cells (**Fig 5A**). Upon OSBP silencing, those cysts which exhibited morphological defects (**Fig 2A**) also showed altered cholesterol distribution. In such cells, the signal of the D4H-cholesterol probe decreased at the plasma membrane and appeared in small, cytoplasmic structures (**Fig 5A**). We found a similar phenotype when we treated MDCK cysts overnight with SWG; however, the drug treatment exhibited a stronger effect as compared to OSBP silencing (**Fig 5B**). Thus, OSBP regulates cholesterol levels of the post-Golgi membranes in polarized epithelial cells.

**Figure 5.**
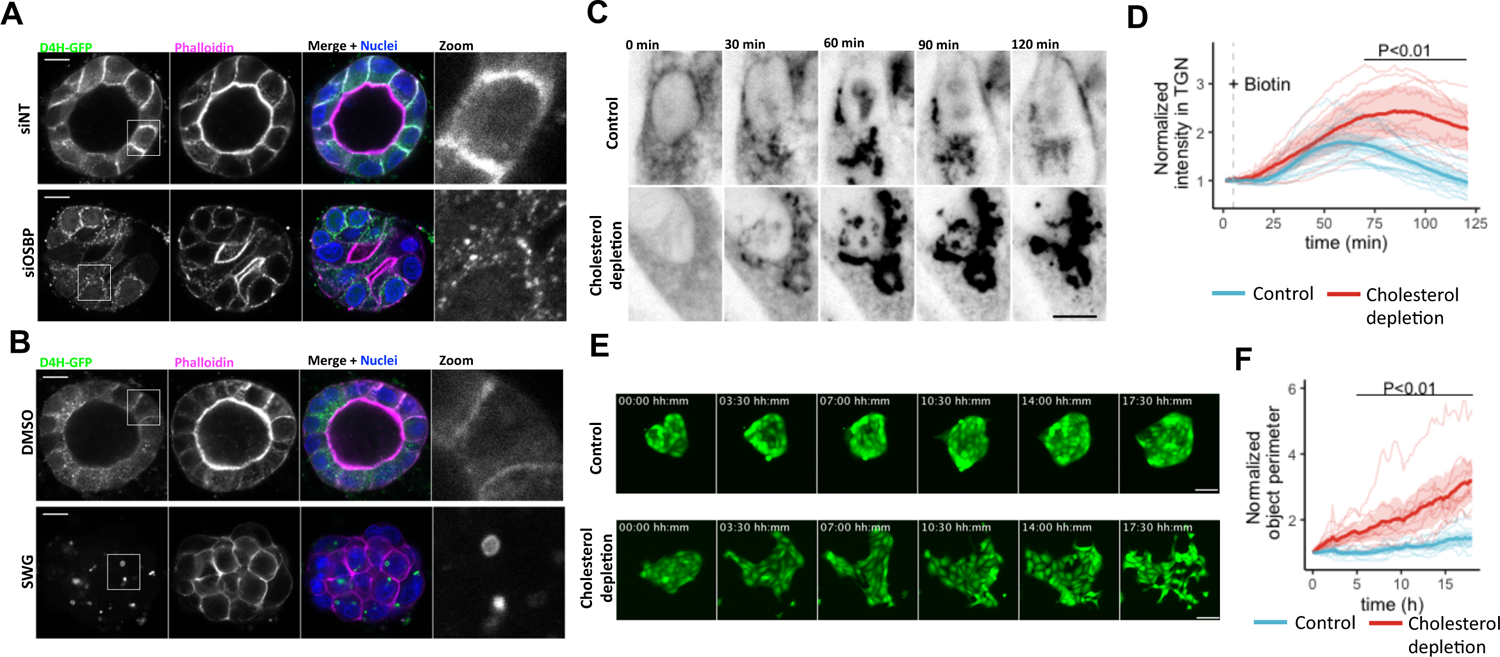
OSBP-derived cholesterol regulates cell polarity and E-cadherin secretion. A. The cholesterol probe D4H-GFP decorates the basolateral membrane of 3D-cultured MDCK cells. Upon OSBP silencing, an altered D4H-GFP distribution can be observed in the cysts displaying morphological defects. Bar=10 #m.
B. Overnight treatment of MDCK cysts expressing the cholesterol probe D4H-GFP results in disrupted distribution of cholesterol. Bar=10 #m.
C. Representative images of live cell microscopy of the effect of cholesterol depletion on E-cadherin secretion. Bar=10 #m
D. Quantification of the live imaging of cholesterol-depleted cells. One representative experiment is shown. Each line corresponds to one cell and bold lines with shaded area indicate mean±95% confidence interval. P-values were calculated by unpaired t test.
E. Representative colonies showing the effect of cholesterol depletion on epithelial cell scattering. Bar=80 #m
F. Cell scattering was quantified by measuring the colony perimeters. Each line corresponds to one colony and bold lines with shaded area indicate mean±95% confidence interval. P-values were calculated by unpaired t test.

To investigate the importance of cholesterol in OSBP-dependent secretion, we followed the OSBP-dependent cargo E-cadherin using the RUSH system in MDCK cells subjected to cholesterol depletion. We found that cholesterol depletion did not affect ER-to-Golgi transition of E-cadherin. However, the Golgi exit of the E-cadherin cargo was blocked in cholesterol depleted cells (**Fig 5C-D, Movie EV4**). Moreover, cholesterol depletion led to scattering of MDCK colonies, further demonstrating the essential function of cholesterol in the secretion of this OSBP-dependent cargo (**Fig 4E, F, Movie EV5**). We concluded that OSBP regulates polarity maintenance and E-cadherin secretion by providing cholesterol to post-Golgi membranes.

### 6. TGFbeta-induced EMT alters cholesterol homeostasis

We found that OSBP is expressed with high level in epithelial cells, while its expression is lower in mesenchymal cells. Since OSBP transfers cholesterol, downregulation of its expression suggests a change in cholesterol homeostasis during EMT, which might contribute to the loss of epithelial characteristics including E-cadherin surface expression. Moreover, epithelial and mesenchymal cells possess different characteristics of cell junction and cell polarity regulation, whose determinants are typically OSBP-dependent secretory cargoes. Therefore, we decided to compare the regulation of the OSBP-dependent cargo secretion in these two cell types.

First, we characterized the changes in sterol homeostasis during TGFbeta-induced mesenchymal progression in the lung adenocarcinoma A549 cells. RT-qPCR data show that genes involved in transcriptional regulation of sterol homeostasis, cholesterol biosynthesis and uptake and efflux, had a decreased expression in TGFbeta-induced mesenchymal cells compared to control cells (**Fig 6A**). Because cholesterol has a preference towards membranes containing saturated lipids, we characterized the acyl-profiles and the relative amounts of phospholipid species as well as ceramides and sphingomyelins in control and TGFbeta-treated cells using mass spectrometry. We calculated a saturation value by dividing the total number of double bonds in the acyl chains by the acyl chain length (number of carbons) and we labelled each lipid species on a blue-red colour gradient, accordingly. Strikingly, the relative quantity of lipids with high unsaturation values (red) increased upon EMT, while the relative level of those characterized by high saturation (blue) showed a decreasing tendency (**Fig 6B, Source Data 2**), indicating that several membrane lipids switch toward unsaturation upon mesenchymal progression.

**Figure 6.**
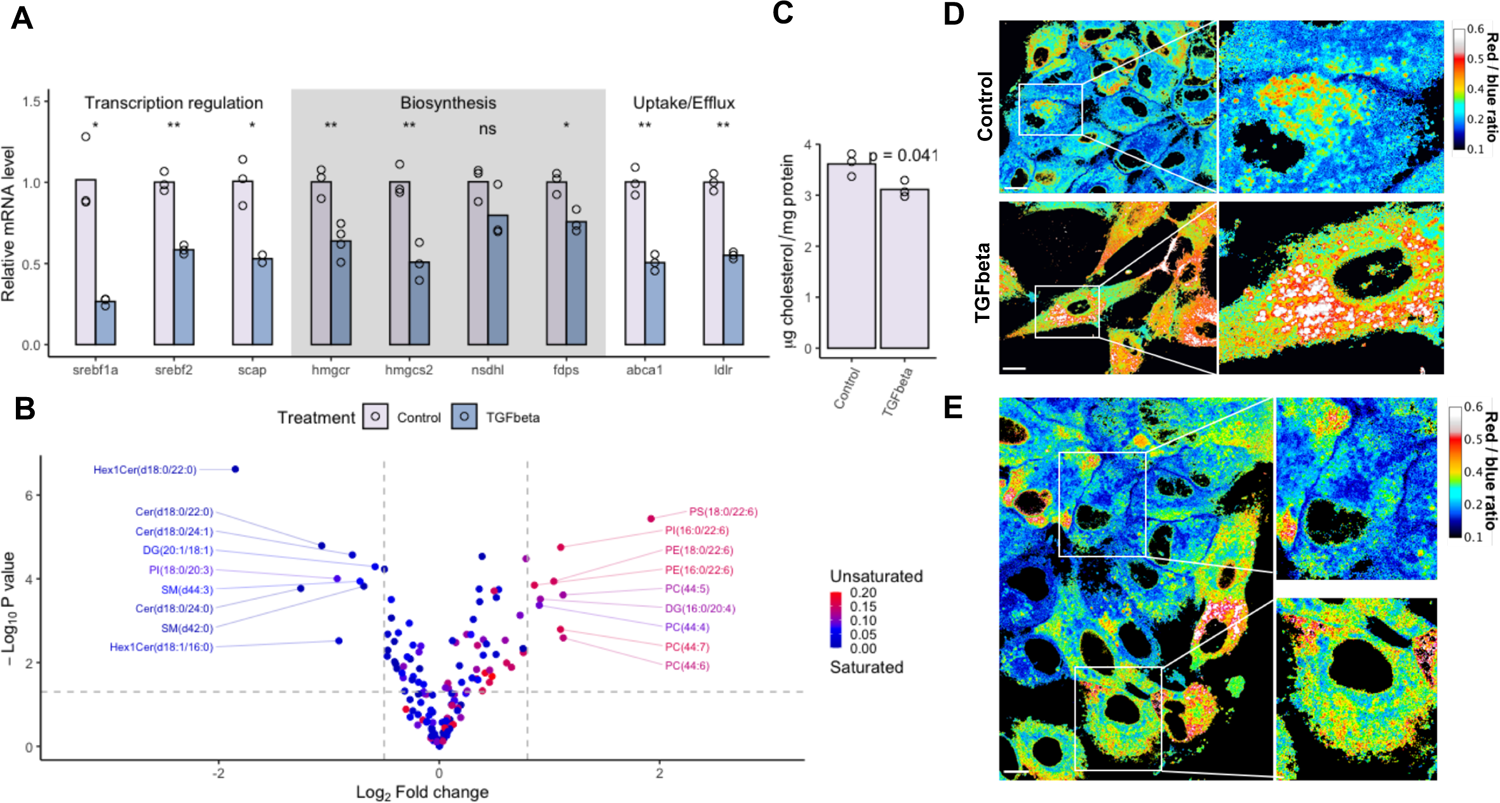
TGFbeta re-wires cholesterol homeostasis. A. Relative mRNA levels of genes involved in cholesterol homeostasis regulation measured by RT-qPCR. Bars show mean of three independent experiments. *p<0.05; **p<0.01; ns=non-significant, Welch’s t test.
B. Volcano plot shows the relative changes of each lipid species detected by mass spectrometry. Color code corresponds to acyl chain saturation. Complete data available in **Source Data 2**.
C. Cholesterol content of control and TGFbeta-treated A549 cells measured by HP-TLC. Bars indicate the mean of three independent experiments. *p<0.05, Welch’s t test.
D. PA-imaging of control and TGFbeta-treated A549 cells shows changes in membrane lipid order upon EMT. Bar=15 #m
E. PA-imaging reveals membrane lipid-order changes upon spontaneous EMT. Bar=15 #m

To test whether this change in acyl profiles was accompanied by a reduction in cholesterol level as well, we performed high-performance thin layer chromatography (HP-TLC) using a cholesterol standard curve to quantify cholesterol in control and TGFbeta-stimulated A549 cells (**Fig EV4A**). As expected, TGFbeta reduced total cholesterol level; TGFbeta-triggered cells contained approximately 14% less cholesterol than the control cells (**Fig 6C**). Changes in other lipid classes such a triglycerides and cholesteryl-esters were also observed, further demonstrating the complex alterations in lipid homeostasis upon EMT (**Fig EV4A-B**).

Finally, we applied the recently developed pyrene-derivative (PA) probe to asses membrane lipid order in living control and TGFbeta-stimulated cells (Niko et al., 2016). As previously reported, cellular membranes show a gradient in the obtained pixel ratio values (Niko et al., 2016). Endomembranes such as Golgi appeared as more polar (red) than the plasma membrane (blue), due to the enrichment of saturated lipids and cholesterol in the PM. Interestingly, PA emission ratio reached the lowest values at cell-cell junctions indicating that these structures favour highly ordered membranes (**Fig 6D**). Strikingly, TGFbeta induced a massive shift in the ratio values toward the red part of the spectrum, indicating that these cells became enriched in disordered membranes, most probably because of lower cholesterol levels and elevated unsaturation (**Fig 6D**).

As an alternative to TGFbeta addition, we scratched an A549 cell layer to promote a spontaneous EMT where cells close to the wound edge migrate toward the cell-free zone. Comparing migrating and non-migrating A549 cells by PA ratiometric imaging showed a comparable phenotype to that observed upon TGFbeta stimulation: migrating cells contain less ordered membranes than their non-migrating counterparts (**Fig 6E**). We concluded that EMT is accompanied by profound changes in bulk lipid homeostasis including cholesterol, which might reflect a decrease in OSBP function and contribute to the loss of epithelial phenotype.

### 7. OSBP-function is downregulated upon EMT

Since cholesterol homeostasis is changed upon TGFbeta treatment, we assessed whether OSBP activity was down-regulated as well during EMT. We started by investigating OSBP expression levels at different stages of EMT. In good agreement with our previous bioinformatics screen and with our RT-qPCR data (**Fig 1**), OSBP protein expression showed a minor time-dependent decrease upon TGFbeta-treatment (**Fig EV5A-B**). We depleted OSBP in A549 cells by siRNA to determine whether OSBP levels affected the expression levels of EMT markers. Despite very efficient OSBP depletion, we observed no changes in EMT marker expression (**Fig EV5C**). Alternatively, we treated cells for 24 h with ORPphilins. Both SWG and OSW-1 treatments lead to the apparition of bands corresponding to proto-E-cadherin, indicating the arrest of its secretion in the TGN (**Fig EV5D**). We found that TGFbeta was able to trigger changes in EMT marker expression upon OSBP silencing, and that TGFbeta-treated A549 cells were able to recover E-cadherin expression after washing out TGFbeta (**Fig EV5E-F**). Thus, OSBP expression itself does not affect the genetic program of EMT, but its inhibition affects the surface targeting of the epithelial determinant E-cadherin.

To achieve efficient lipid transfer, OSBP constantly cycles between the cytosol and Golgi membranes, thus when it is highly active, it shows a lower Golgi recruitment in fixed cells. In contrast, less cycling between its membrane bound and cytosolic form is coupled to a lower lipid transfer activity, which leads to higher TGN-recruitment of OSBP due to the elevated level of PI(4)P in the Golgi membranes (Mesmin et al., 2013). In untreated A549 cells, OSBP was detected both in the cytoplasm and in the *trans*-Golgi network labelled by anti-TGN46. In TGFbeta-induced mesenchymal cells, OSBP accumulated in the *trans*-Golgi at the expense of its cytoplasmic pool (**Fig 7A-B**). We observed a similar trend in MDCK cells upon treatment with HGF, which is a more potent EMT-inducer than TGFbeta in this cell line (**Fig EV5G, H**) (Howard et al., 2011).

**Figure 7.**
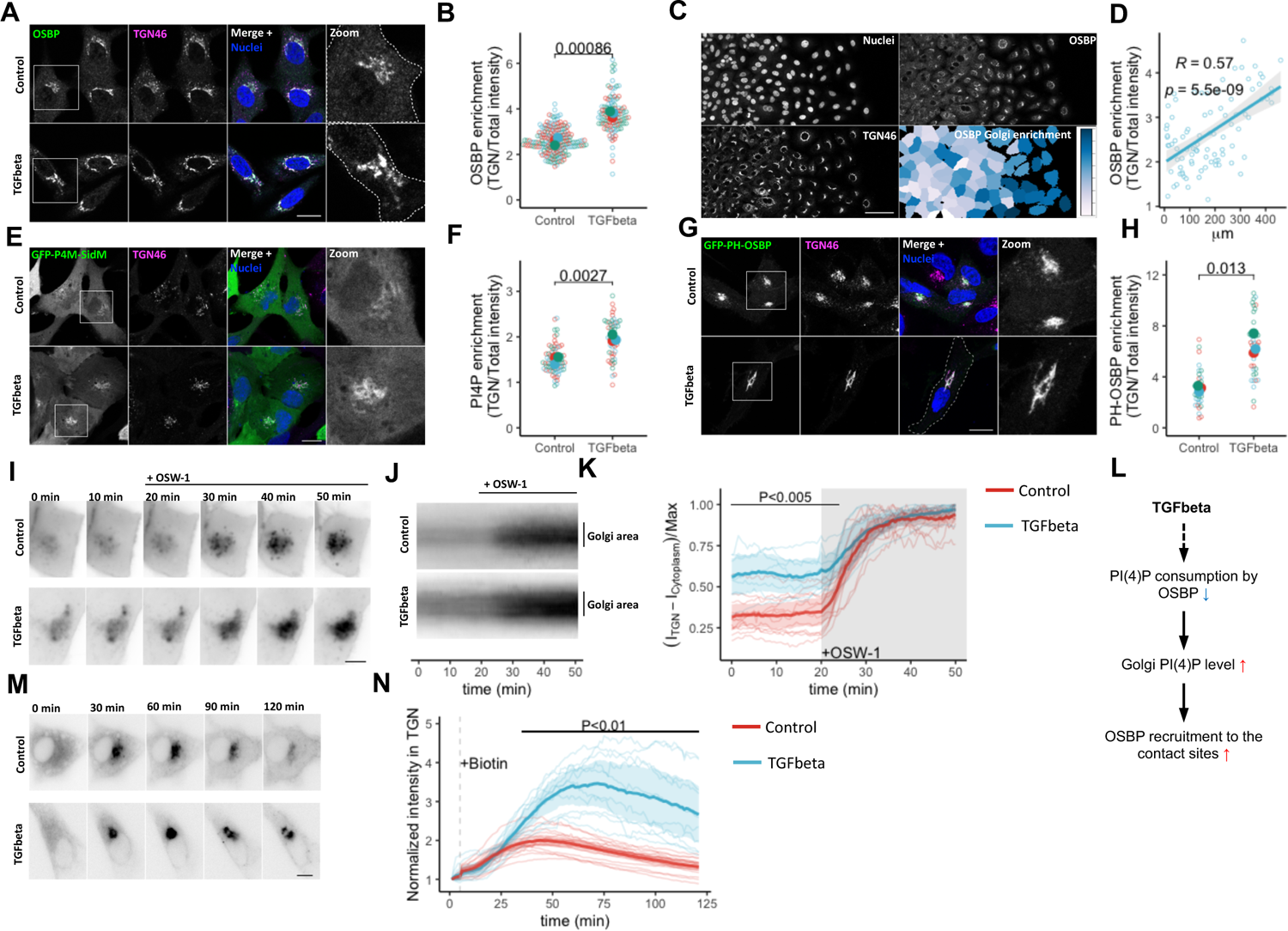
Mesenchymal progression is associated to changes in OSBP functions. A. Confocal microscopy shows the re-arrangement of OSBP pool in control and TGFbeta-treated A549 cells. Bar=20 #m.
B. Superplot shows the enrichment of OSBP signal in TGN in control and TGFbeta-treated cells. Means obtained from three independent experiments were compared by Welch’s t test.
C. Changes in TGN-enrichment of OSBP signal upon wound healing of A549 cells. Pseudo-colored image indicates TGN localised /Total OSBP intensity ratio. Bar= 80 #m.
D. OSBP TGN enrichment values upon wound healing. Each point represents one cell. Values on the x axis indicate the distance of the cells from the left border of the image.
E. Confocal images of control and TGFbeta-treated A549 cells expressing the PI(4)P reporter GFP-P4M-SidM. Bar=15 #m.
F. Superplot shows the enrichment of GFP-P4M-SidM signal in TGN in control and TGFbeta-treated cells. Means obtained from three independent experiments were compared by Welch’s t test.
G. Confocal images of control and TGFbeta-stimulated A549 cells expressing GFP-PH-OSBP. Bar=15 #m.
H. Superplot shows the enrichment of GFP-PH-OSBP signal in TGN in control and TGFbeta-treated cells. Means obtained from three independent experiments were compared by Welch’s t test.
I. Live imaging of control and TGFbeta-treated A549 cells expressing GFP-P4M-SidM. 50nM OSW-1 was added to the cells 20 min after the start of the imaging. Bar=10 #m
J. Averaged kymographs of Golgi regions of control and TGFbeta-treated A549 cells. Control n= 12, TGFbeta n=8. P-values were calculated by unpaired t test.
K. Kinetics of the GFP-P4M-SidM experiments. One representative experiment is shown. Each line corresponds to one cell and bold lines with shaded area indicate mean±95% confidence interval. P-values were calculated by unpaired t test.
L. Schematic representation of how TGFb-triggered EMT leads to elevated Golgi accumulation of OSBP.
M. Live epifluorescent imaging of control and TGFbeta-treated cells shows different kinetics of E-cadherin secretion. Bar=10 #m
N. Quantification of the live imaging. One representative experiment is shown. Each line corresponds to one cell and bold lines with shaded area indicate mean±95% confidence interval. P-values were calculated by unpaired t test.

To examine whether this redistribution of OSBP could be observed when epithelial cells undergo EMT spontaneously, we cultured cells until they reached confluency, followed by scratching the cell layer to obtain a cell-free area. In the following 24 hours, cells in the proximity of the wound edge migrated into the cell-free zone and underwent spontaneous EMT, while the remaining cells preserved their epithelial features. The immunofluorescent signal corresponding to endogenous OSBP showed a gradual *trans*-Golgi enrichment in the cells closer to the wound edge (**Fig 7C-D**), confirming a correlation between EMT and a shift in the partitioning of OSBP between the cytosol and ER-trans Golgi membrane contact sites. The consistent shift of OSBP towards the TGN upon various EMT-triggering approaches suggests downregulation of its lipid transfer activity.

Since OSBP is recruited to the contact sites via its PI(4)P-binding PH-domain, an elevated Golgi PI(4)P level could explain its increased recruitment to the Golgi membranes in mesenchymal cells. In agreement with this, it has been suggested that mesenchymal cells have higher PI(4)P levels compared to epithelial cells (Tokuda et al., 2014). To measure the level of PI(4)P at the Golgi, we ectopically expressed the PI(4)P reporter GFP-P4M-SidM in control and in TGFbeta-treated A549 cells, followed by labelling the *trans*-Golgi Network with a TGN46-specific antibody (**Fig 7E**). Quantification of the confocal micrographs shows that the level of PI(4)P at the Golgi increased in TGFbeta-triggered mesenchymal cells (**Fig 7F**). A similar phenotype was observed when we compared the localization of a GFP-fused form of the PI(4)P binding PH-domain of OSBP (GFP-PH-OSBP) in epithelial and mesenchymal cells confirming that the increase in OSBP at the TGN is due to elevated PI(4)P binding (**Fig 7G-H**).

We previously reported that about half of the total cellular PI(4)P pool is consumed by OSBP (Mesmin et al., 2017). Therefore, an elevated Golgi PI(4)P level might reflect that OSBP is biochemically less active in the contact sites. To determine whether the difference in PI(4)P levels between epithelial and mesenchymal cells was due to differences in the biochemical activity of OSBP, we measured the increase in PI(4)P at the TGN upon OSW-1 addition. By inhibiting OSBP, this drug protects PI(4)P from the OSBP lipid exchange cycle; thus, comparing steady-state and maximal values allows the estimation of the pool of PI(4)P that is consumed by OSBP. Following OSW-1 addition, Golgi-specific PI(4)P levels increased rapidly and then reached a plateau within 15 min in both epithelial and mesenchymal cells (**Fig 7I-K**). However, the difference between the steady state and the maximal value was bigger in epithelial cells than in mesenchymal cells (**Fig 7K**). Quantification of the PI(4)P signal before and after OSW-1 addition suggested that OSBP consumed ca. 70% of the total cellular PI(4)P pool in control A549 cells compared to only ∼40% in TGFbeta-treated cells, indicating a significant reduction of OSBP activity between control and TGFbeta-stimulated cells (**Fig 7K**). Performing a similar experiment on HGF-stimulated MDCK cells stably expressing the GFP-P4M-SidM sensor led to similar results (**Fig EV5I-J**). Altogether, these results indicate that OSBP is biochemically less active in TGFbeta-triggered mesenchymal cells, and that the excess of Golgi-localised PI(4)P in such cells results in the elevated recruitment of OSBP to the contact sites (**Figure 7L**).

Considering the importance of OSBP in the traffic of E-cadherin and the diminished OSBP activity upon mesenchymal progression, we finally asked whether forced secretion of this OSBP-dependent cargo in TGFbeta-stimulated cells followed the same kinetics and trafficking efficiency as in their non-stimulated counterparts. For this, we expressed GFP-tagged E-cadherin as a RUSH cargo in control and in TGFbeta-induced mesenchymal cells. Live cell imaging experiments showed that mesenchymal cells accumulated much more E-cadherin in the Golgi due to delayed exit of E-cadherin from Golgi-structures as compared to control cells (**Fig 7M-N, Movie EV6**). Altogether, these experiments suggest that upon EMT, cells downregulate not only the expression of the epithelial proteins but also reduce OSBP-dependent lipid exchange, which assures the subcellular distribution of epithelial determinants such as E-cadherin.

## Discussion

Among the members of the ORP protein family, we identified OSBP as a prominent epithelial cell-specific gene. Moreover, we found that OSBP inhibition by the specific drug SWG lead to the loss of epithelial polarity, demonstrating a key function of OSBP in maintaining the epithelial phenotype. In addition, cysts developed from OSBP-silenced cells also displayed higher percentage of morphological defects than their controls. Because OSBP transfers cholesterol and PI(4)P at ER-TGN contact sites, we hypothesised that by regulating Golgi lipid environment, OSBP controls cargo-specific secretion to regulate epithelial polarity.

By exploiting a TurboID-based proximity ligation, we detected numerous plasma membrane-localised proteins and their cytoplasmic associated partners in the vicinity of OSBP. The repertoire of OSBP partners includes many components of cell junction complexes such as adherent junctions, tight junctions, desmosomes, focal adhesions and gap junctions, which are key determinants of epithelial polarity. Upon OSBP inhibition, the majority of the cell junction and adhesion molecules become enriched in the OSBP proximity proteome. Since this condition also enriches OSBP at the ER-TGN contact sites, we suggest that these cargoes go to the proximity of such contact sites when they are subjected to Golgi secretion.

By combining live cell imaging and synchronized cargo release from the ER, we discovered that the secretion of one of the identified OSBP proximity proteins, E-cadherin, is arrested in the Golgi upon OSBP inhibition, showing that OSBP activity is necessary for post-Golgi E-cadherin secretion and surface expression. We found that, similarly to E-cadherin, endogenous beta-catenin distribution is also affected by OSBP inhibition, indicating that non-transmembrane components of the adherent junctions also depend on OSBP for their trafficking. Because adherent junction complexes are assembled at least partially during Golgi maturation, it is not surprising that they move together along the late secretory pathway, thereby being affected *en bloc* by OSBP inhibition (Chen et al., 1999; Curtis et al., 2008; Wahl et al., 2003).

Importantly, depleting cholesterol from the membranes phenocopied the effects of ORPphilin treatments on E-cadherin secretion, indicating that OSBP-provided cholesterol is the key molecule of OSBP-regulated secretion. Because cadherin molecules are embedded in cholesterol-rich plasma membrane regions (Guillaume et al., 2013; Resnik et al., 2019; Seveau et al., 2004), we propose that this membrane environment is assembled upon Golgi sorting by the contribution of OSBP-mediated cholesterol transfer and is a pre-requisite for efficient *trans-* Golgi trafficking of E-cadherin. Notably, a class of TGN-derived transport carriers named CARTS (<u>Car</u>riers of the <u>T</u>GN to the cell <u>s</u>urface) have been already identified as cholesterol dependent transport vesicles (David & Castro, 2021; Wakana et al., 2021). Although the purification of this vesicle fraction is demanding, early proteomic characterization identified a high number of desmosomal proteins in such transport vesicles (Wakana et al., 2012). As weidentified major desmosomal proteins in the OSBP proximity proteome, an overlap between CARTS cargoes and OSBP-dependent secretory clients is likely. It has long been suggested that cholesterol-enriched microdomains in the TGN facilitate the apical secretion of some cargoes (Cao et al., 2012; Keller & Simons, 1997). However, we identified both basolateral and apical cargoes in the OSBP-proximity ligation assay. Thus, we propose that OSBP-derived cholesterol facilitates cargo clustering in the TGN membrane to distribute surface proteins of epithelial cells. How this assembly process, which should require spatio-temporal coordination between ER-TGN contact sites and the formation of trafficking intermediates at the TGN precisely occurs, will require further investigation. We note that the contact sites involving OSBP are highly dynamic (Mesmin et al., 2017) and engage only a fraction of the TGN membranes, which might favour the formation of transport intermediates in their close proximity.

When cells undergo a phenotypic switch such as differentiation, they must not only change their overall proteomic profiles, but also re-shape their secretory routes to better support the trafficking of their actual cargo set. The epithelial-to-mesenchymal transition is an ideal model to observe such changes for multiple reasons. First, expression of cell-cell junction components is reduced in mesenchymal cells to prime the individualization of the cells. Therefore, many of the identified cargoes of the OSBP-regulated secretion are downregulated in these cells. Second, similar to others, we observed major changes in membrane lipid composition upon EMT, which is coupled to re-wiring of cholesterol homeostasis (Giudetti et al., 2019; Sampaio et al., 2011). Since we previously found that OSBP consumes less PI(4)P when the level of cholesterol to be transported is lower, the altered cholesterol homeostasis suggests adaptation of the OSBP cycle upon changes in EMT status as well (Mesmin et al., 2017). Third, our initial meta-screen on RNAseq data shows that OSBP is predominantly expressed in cells with high epithelial scores, while in mesenchymal cells, its expression is lower, suggesting that there is a demand for elevated OSBP function in epithelial cells. In addition to this, cohorts indicate that low OSBP expression is coupled to poor patient survival in some cancers, which further suggests that high OSBP levels are associated with a more differentiated – therefore less fatal – cellular phenotype.

We found that OSBP burns less PI(4)P in mesenchymal cells than in epithelial cells, indicating that its lipid transfer activity is downregulated upon EMT. Furthermore, we found that the mesenchymal Golgi environment is less permissive to process high cholesterol-demanding model cargoes such as E-cadherin. These findings highlight a new aspect of the EMT program: a decrease in OSBP activity and OSBP-dependent cargo secretion. To achieve this, the cell might control OSBP at three levels: (i), downregulate its protein expression, (ii), cut off cholesterol flux from OSBP, (iii), regulate its activity by posttranslational modifications e.g. phosphorylation (Goto et al., 2012). Notably, we found that that OSBP protein level is slightly decreased upon EMT, and we detected reduction of total cholesterol content upon EMT as well. Whether these changes are the determining reasons for the 2-fold decrease in OSBP-dependent PI(4)P consumption upon EMT remains to be elucidated.

OSBP silencing by siRNA and OSBP blocking by ORPphilins do not necessarily lead to the same phenotype. For example, SWG treatment has a massive effect on cholesterol distribution and cyst morphology, while OSBP silencing triggers similar, but less prominent effects. ORPphilins stabilizes OSBP at ER-TGN contact sites, thereby freezing its dynamics. In contrast, silencing OSBP expression reduces its level at the contact sites, which might let other ORPs compensate for the lack of OSBP function, thus favouring rescue phenotypes, an effect previously observed for the related Osh family in yeast (Beh et al., 2001). Burgett et al. also found that silencing of OSBP expression does not precisely phenocopy ORPphilin treatments on cell proliferation. Whether this is due to the activation of compensatory mechanisms, needs to be studied. Nevertheless, these differences highlight the importance of having specific pharmacological tools to study OSBP functions in a spatiotemporal manner in cells.

Among the OSBP proximity proteins, we identified a set of cytoplasm-localised factors, which show a tendency to be enriched in the proximity of the non-membrane bound, soluble form of OSBP. OSBP has already been reported to function as a cytoplasmic scaffold protein to initiate the assembly of phosphatase-containing complexes (Wang et al., 2005). In good agreement with this finding, we identified numerous phosphatases and kinases in the OSBP proximity landscape, suggesting that the function of OSBP might not be limited to cholesterol transport but extend to cell signalling.

In conclusion, our work contributes to a better understanding of how lipid transfer proteins regulate secretory mechanisms of specific cargoes. The OSBP-dependent secretory pathway shows high cell environment-dependent plasticity and is regulated upon EMT. Since EMT is considered to be a pre-requisite for metastasis and is a hallmark of cancer malignancy, analysing the OSBP-dependent secretory pathway, should help understanding malignant secretion and serve to identify new targets for rational therapeutic developments.

## Materials and Methods

### Plasmids and reagents

To generate OSBP-TurboID-HA, we amplified OSBP cDNA using sequence specific primers containing NheI and HindIII cutting sites from an OSBP-mCherry expressing plasmid described earlier (Mesmin et al., 2013), then the amplicon was inserted into a pcDNA3.1-Zeo(+) vector. Using specific primers containing HindIII and BamHI restriction sites TurboID was amplified from a reference template plasmid and the sequence was inserted into this vector to generate an OSBP-TurboID-HA expressing construct. In order to replace the EGFP-tag with mCherry in the Str-KDEL_SBP-EGFP-Ecadherin construct, EGFP was digested from the plasmid using SbfI and FspI restriction enzymes then the mCherry cDNA amplified by sequence-specific primers was inserted into the plasmid by blunt ligation. All the generated plasmids were subjected to capillary sequencing prior to application to confirm that their sequences were correct. For detailed information on origin, application and reference of plasmids and reagents used in this study, see **Supplementary Table**.

### Cell culture, EMT, plasmid and siRNA transfection, 3D cyst formation

MDCK cells were cultured in Minimal Essential Medium with Earl’s balance salts (EMEM, Sigma-Aldrich) completed with 5% foetal bovine serum and 1% ZellShield (Minerva Biolabs). A549 cells were maintained in Dulbecco’s Modified Eeagle’s Medium (DMEM, Gibco) complemented with 10% FBS and 1% ZellShield. Cells were maintained in a 37°C humidified incubator kept in a 5% CO_2_ atmosphere. To transfect plasmid DNA and siRNA (Thermo Fisher Scientific), MDCK cells were nucleofected by Amaxa Cell Line Nucleofector using Kit L (Lonza). Plasmid DNA was transfected into A549 cells by Xfect transfection reagent (Takara Bio), while transfection of ON-Targetplus siRNAs (Dharmacon) was carried out by Lipofectamine RNAiMAX Transfection Reagent (Invitrogen) following the instructions of the manufacturer. siRNA target sequences can be found in **Supplementary Table**. To induce EMT in A549 cells, 0.25 x 10^6^ cells were seeded into 10 mL cell culture dishes then treated with 5 ng/mL TGFbeta (Abcam) on the next day. TGFbeta was kept on the cells for at least 72 h or for indicated times. Cells were re-seeded to appropriate culture dishes for experiments, keeping a 5 ng/mL TGFbeta concentration in the medium throughout the experiment. A similar approach was used for MDCK cells, however, these cells were stimulated with 40 ng/mL HGF for 24 h. To form MDCK cysts, 2000 trypsin-individualised cells were re-suspended in 20 μL Matrigel (Cornig) and dropped on 8 or 4 well μ-Slide chambered coverslips (Ibidi). Following the polymerization of the gel, cell culture medium was placed on the samples and cysts were let to develop for 72 h.

### Generation of stable cell lines

To generate stable cell lines, MDCK cells were transfected with linearized plasmid DNA and selected with either 500 μg/mL G418 (Sigma-Aldrich) or 400 μg/mL Zeocin (Gibco) 48 h following transfection for 2 weeks. Surviving colonies were picked and tested for transgene expression by immunoblotting or fluorescent microscopy. For the proximity labelling assays, clones expressing OSBP-TurboID-HA with a similar level to that of the endogenous OSBP were selected.

### RT-qPCR

To analyse mRNA expression of selected genes, 0.2×10^6^ A549 cells were seeded to 6 cm diameter culture dishes, then EMT was induced with 5 ng/mL TGFbeta for 72 h. Total RNA was purified using RNeasy Plus Mini kit (Qiagen), and a total amount of 2 μg RNA was reverse transcribed with a GO Script Reverse Transcriptase kit (Promega) using random primers. Relative transcript levels were measured in a LightCycler^®^ 480 Instrument (Roche) using Takyion^TM^ No Rox SYBR^®^ MasterMix (Eurogentec). Gene-specific primer sequences are summarized in **Supplementary Table**. To calculate gene expression levels, the ΔΔCT method was applied using GAPDH as a reference gene.

### Immunofluorescence and confocal microscopy

For immunofluorescent labelling, cells were seeded into 8 or 4 well μ-Slide chambered coverslips (Ibidi). Following the experimental procedures, cells were fixed using 4% PFA and quenched in 50 mM NH_4_Cl. Cells were permeabilized with 0.5% Saponin in 2% BSA-PBS then antibodies were diluted in the same buffer and incubated with the cells overnight at 4°C. Following three washing steps with PBS, secondary antibodies were applied for 1 h together with Hoechst (Invitrogen) and other dyes such as DraQ5 (Cell Signaling Technology), Phalloidin (Invitrogen) and Streptavidin594 (Invitrogen) depending on the experiment. Detailed list of antibodies and other materials applied for immunolabelling is collected in **Supplementary Table**.

Immunolabelled samples were analysed with a Zeiss LSM 780 confocal microscope operated with ZEN software using a Plan-Apo-chromat 63X/1.4 Oil objective (Carl Zeiss). CellProfiler^TM^ 4.1.3 image analyses software (Mcquin et al., 2018) was applied to assess confocal micrographs by custom-made pipelines.

### Epifluorescent live cell imaging and image analysis

For live epifluorescent imaging experiments, nucleofected MDCK cells were seeded into 35 mm diameter μ-Dishes (Ibidi) in 5 x 10^5^ cell/dish density and incubated 24 h before imaging. Control or TGFbeta-stimulated A549 cells were seeded at 0.2 x 10^6^. Prior to imaging, cells were washed three times with pre-warmed PBS and medium was replaced with 1 mL phenol red-free culture medium and samples were mounted in a 37°C stage chamber (Okolab). For RUSH experiments, release of secretory model cargoes was induced by adding biotin to the cells at 80 μM with DMSO or ORPhilins at the indicated time points. Time-lapse imaging was carried out by a IX83 inverted microscope (Olympus) equipped with an iXon3 camera (Andor) and an UPlanSApo 60X/1.35 oil objective (Olympus) and operated by a MetaMorph software (Molecular Devices). Generated image sequences were analysed by Fiji image processing software (Schindelin et al., 2012) using StackReg plugin. Data analyses and plotting was carried out in Rstudio using ggplot2.

### Proximity labelling and proteomics analyses by NanoHPLC-HRMS

To map the OSBP proximity interactome, 1 x 10^7^ wild type MDCK or MDCK cells stably expressing OSBP-TurboID were seeded into 150 mm diameter dishes in triplicates/condition. The next day, cells were either treated with DMSO or with 1 μM SWG. After 50 min, biotin was added to the culture mediums at 0.5 mM concentration and samples were incubated for an additional 10 min at 37°C to complete a 1 h SWG treatment time. Cells were then washed 5 times with ice-cold PBS and lysed in RIPA buffer supplemented with cOmplete Protease Inhibitor Cocktail (Roche). Cell lysis was facilitated by sonication and following centrifugation, protein concentrations were determined from the supernatant using a BCA assay kit (Thermo Fisher Scientific). Lysates were subjected to immunoblotting to confirm OSBP-TurboID expression and successful protein biotinylation (**Figure EV2C**). Sample preparation for mass spectrometry was carried out as described elsewhere (Branon et al., 2018; Hung et al., 2016) with minor modifications. Briefly, 5 mg of total protein/sample was combined with 330 μL RIPA-equilibrated Pierce^TM^ Streptavidin Magnetic Beads (Thermo Fisher Scientific) and incubated on a rotator at 4°C overnight. Beads were washed two times with RIPA buffer, then once with 1 M KCl, 0.1 M Na_2_CO_3_, with 2 M urea diluted in 10 mM TRIS-HCl pH=8.0 and twice with RIPA buffer again. Approximately 5% of the slurry was saved to verify the successful enrichment of biotinylated material by silver staining (**Figure EV2D**). Beads were further washed twice with 50 mM Tris-HCl pH=7.5 and twice with 2 M urea diluted in 50 mM Tris-HCl pH=7.5 then on-bead digestion was performed by adding 0.4 μg Pierce^TM^ MS grade trypsin protease (Thermo Fisher Scientific) and 1 mM DTT in 2 M urea/50 mM Tris-HCl pH=7.5. Trypsinization was performed at 25°C with agitation for 1 h, supernatant was removed and beads were washed two times with 2 M urea/50 mM TRIS-HCl pH=7.5, then the supernatant was combined with the washes. Peptides were reduced with 4 mM DTT for 30 min at 25°C and alkylated by 10 mM iodoacetamide (Sigma-Aldrich) for 45 min in dark in a thermoshaker. To complete digestion, 0.5 μg trypsin was added to each sample and digested overnight at 25°C. Following this, peptides were desalted on Pierce^TM^ Peptide Desalting Spin Columns (Thermo Fisher Scientific) and concentrated in a SpeedVac concentrator.

NanoHPLC-HRMS analysis was performed using a nanoRSLC system (ultimate 3000, Thermo Fisher Scientific) coupled to an Easy Exploris 480 (Thermo Fisher Scientific). Peptide separation was carried out using the Easy-nLC ultra high performance LC system. 5 µL of peptide solution was injected and concentrated on a µ-Precolumn Cartridge Acclaim PepMap 100 C18 (i.d. 5 mm, 5 mm, 100 Å, Thermo Fisher Scientific) at a flow rate of 10 mL/min and using solvent containing H_2_O/ACN/TFA 98%/2%/0.1%. Next peptide separation was performed on a 75 mm i.d. x 500 mm (2 µm, 100 Å) PepMap RSLC C18 column (Thermo Fisher Scientific) at a flow rate of 300 nL/min. Solvent systems were: (A) 100% water, 0.1%FA, (B) 100% acetonitrile, 0.1% FA. The following gradient was used t = 0 min 2% B; t = 3 min 2%B; t = 103 min, 20% B; t = 123 min, 32% B; t = 125 min 90% B; t = 130 min 90% B; (temperature was regulated at 40°C). MS spectra were acquired at a resolution of 120 000 (200 m/z) in a mass range of 375–1500 m/z with an AGC target 3e6 value of and a maximum injection time of 25 ms. The 20 most intense precursor ions were selected and isolated with a window of 2 m/z and fragmented by HCD (Higher energy C-Trap Dissociation) with a normalised collision energy (NCE) of 30%. MS/MS spectra were acquired in the ion trap with an AGC target 5e5 value, the resolution was set at 15,000 at 200 m/z combined with an injection time of 22 ms.

### Mass spectrometry data analyses

MS data were subjected to LFQ analysis using MaxQuant v1.6.17.0 (http://www.maxquant.org/) using the MaxQuant platform (v1.6.7.0). Database search of the MS/MS data was performed in MaxQuant using the Andromeda search engine against *Canis lupus familiaris* UniProtKB database (Nov. 2020) and MaxQuant contaminants database. Digestion mode was set to Trypsin/ P specificity, with a fixed carbamidomethyl modification of cysteine, and variable modifications of protein N-terminal acetylation and methionine oxidation. Mass deviation was set to 20 ppm for the first and 6 ppm for the main search and the maximum number of missed cleavages was set to 2. Peptide and site false discovery rate (FDR) were set to 0.01, respectively. The search for co-fragmented peptides in the MS/MS spectra was enabled (“second peptides” option). Protein identification was performed using a minimum of 2 unique and razor peptides. Quantification was achieved using the LFQ (Label-Free Quantification) algorithm. Razor and unique peptides were used for LFQ quantification and the “label min. ratio count” was fixed at 2. The match between runs option was enabled, allowing a time window of 0.7 min to search for already identified peptides in all obtained chromatograms. Raw intensity values corresponding each identified proteins were used to calculate Log_2_Fold change and -Log_10_P values using Welch’s t-test. Identified proteins were subjected to enrichment analyses by the online tool www.metascape.org (Zhou et al., 2019) using selected functional background gene sets with a 0.01 P-value cutoff, while Panther GO tool (www.pantherdb.org) (Thomas, 2003) was used for functional annotation.

### Lipidomics analysis

Briefly, a modified Bligh and Dyer (Bligh & Dyer, 1959) extraction was carried out on cell pellets in order to extract lipids, which were then separated by chromatography with a C18 column and an appropriate gradient of mobile phase. The mass spectrometry analyses were done using a Q-Exactive mass spectrometer (Thermo Fisher Scientific) operating in data dependent MS/MS mode (dd-MS2) and the data were then processed using LipidSearch software v4.2.21 (Thermo Fisher Scientific) in product search mode. Detailed protocol for lipidomics analyses is included in the supplementary methods.

### Immunoblotting

For immunoblotting, whole cell lysates were obtained by lysing the cells in RIPA buffer containing cOmplete Protease Inhibitor Cocktail (Roche). Following sonication, samples were pelleted and supernatant was used as whole cell extracts. Protein concentration of each sample was determined as described before and a total amount of 20 μg protein/sample was resolved on 4-20% gradient gels (Bio-Rad). Proteins were transferred onto nitrocellulose or PVDF membranes futher blocked with 5% non-fat dry milk-TBST. To reveal biotinylated proteins, the membranes were probed with HRP-conjugated Streptavidin-HRP diluted in 3% BSA-TBST. For antibody reactions, samples were transferred to PVDF. Following blocking, primary antibodies were diluted in 1% non-fat dry milk-TBST and incubated with the membranes overnight at 4°C. HRP-conjugated secondary antibodies were applied then signal was developed by a chemiluminescent HRP substrate (Millipore) and detected with a Fusion FX7 instrument (Vilber Loumat). Antibodies and their respective working dilutions used for immunoblotting are summarised in **Supplementary Table**.

### Scattering assay and image analysis

For cell scattering assays, MDCK-EGFP cells were seeded into 12 well plates at 2000 cell/well density, then cell clusters were left to grow for 3 days. Following the addition of ORPhilins or cholesterol depletion, time-lapse imaging was performed in a Cytation5 Cell imaging multimode-reader set to 37°C, containing 5% CO_2_ and operated by a Gen5 software. Cluster scattering was quantified by measuring object perimeters through the time-laps imaging by CellProfiler^TM^ 4.1.3 image analyses software.

### Pharmacology and cholesterol depletion

OSW-1 was a generous gift from Matthew D. Shair (Harvard) while SWG was provided by Fanny Roussi (CNRS ICSN). For the proximity biotinylation assay, SWG was used at 0.5 μM concentration for 1 h. Upon RUSH experiments, OSW-1 was applied at 20 nM, while SWG was used at 0.5 μM. SWG was used overnight on MDCK cysts at 0.5 μM. For experiments shown Figure 4d, e and h, OSW-1 was applied at 2 nM, while SWG was used at 0.5 μM. To measure OSBP-dependent PI(4)P consumption in living cells, OSW-1 was applied at 50 nM on GFP-P4M-SidM expressing cells. For cholesterol depletion of MDCK cells, samples were treated with 5 mM Methyl-β-cyclodextrin (Sigma-Aldrich) diluted in 10 μM Lovastatin (Sigma-Aldrich) and 5% lipoprotein-depleted serum-containing medium for two hours, then after 5 washes with PBS, cells were incubated in 10 μM Lovastatin and 5% lipoprotein-depleted serum-containing medium throughout the experiments.

### Lipid order imaging

The lipid order probe PA was a kind gift from Andrey S Klymchenko (CNRS, Illkirch). To analyse lipid order of living cells, control and TGFbeta-stimulated cells were washed three times with pre-warmed PBS and PA dye was incubated with the cells for 10 min in phenol red-free cell culture medium. Following incubation, cells were imaged at 37°C by a Zeiss LSM 780 confocal microscope operated with ZEN software. Samples were excited by 405 nm and fluorescence in two emission windows (470-550 nm and 550-700 nm) was detected. Ratiometric images were generated using Fiji software by calculating ratio values between long and short wavelength channels, then the resulted images were converted to pseudocolor scale.

### Bioinformatics, data analyses and statistics

For correlation analyses, we extracted EMT marker as well as ORP expression data from publicity available TCGA data using www.cbioportal.org (Cerami et al., 2012). For ORP expression analyses, a similar approach was used which was described elsewhere (Rajapakse et al., 2018). For cell line expression data, we used The Cancer Cell Line Encyclopedia set (Barretina et al., 2012), and a TCGA dataset was used to analyse lung adenocarcinoma patient data (Hoadley et al., 2018). A previously identified EMT signature gene set was used to define EMT characteristics (Kohn et al., 2014). To calculate the epithelial-mesenchymal index the following method was applied: first, mean expression values for epithelial and mesenchymal markers were obtained (Mean_mes_ and Mean_epi_), then a cell line/sample-specific EMT value was calculated by subtracting Mean_epi_ from Mean_mes_ and multiplying the value by −1. This formula was used to calculate a value for all cell lines/tumour samples, then each value was normalised to the mean EMT value of the total data set. Data visualization and correlation analyses were carried out using ggplot2 with GGally extension and heatmap.plus package in Rstudio.

To obtain publicly available survival data we used the online tool www.kmplot.com (Győrffy, 2021). Briefly, expression data for OSBP and OSBPL2 was analysed in 719 patient samples identified histologically as lung adenocarcinoma. Patients were split by median expression to obtain high and low expression categories. For data visualisation and data analyses we used ggsurvplot function in ggplot2 using Rstudio. To meta-analyse histology data we used the human proteome atlas (Uhlen et al., 2015).

Raw data analyses and plotting were carried out in Rstudio using ggplot2 and other appropriate packages. Superplots were used to show data obtained from repeated experiments and were prepared according to the guidelines published elsewhere (Lord et al., 2020). On these plots, each colour represents data obtained from one experiment, and filled dots correspond to mean values. Mean values between conditions were compared by unpaired t-test to confirm statistical significance. Differences were considered statistically significant when p values were smaller than 0.05. Other statistical tests used for data analyses are indicated in each figure legend.

**Supplementary methods**

## Chemicals used for lipid analyses

Extraction: 2,6-Di-tert-butyl-4-methylphenol or [(CH_3_)_3_C]_2_C_6_H_2_(CH_3_)OH (Sigma, 34750), Chloroform or CHCl_3_ (Sigma, 34854), Methanol or CH_3_OH (Sigma, 34860).

Quantification: Sodium dihydrogen phosphate dihydrate or NaH_2_PO_4_·2H_2_O (VWR, 28011.291), Ammonium molybdate tetrahydrate or (NH_4_)_6_Mo_7_O_24_·4H_2_O (Sigma, A7302), L-ascorbic acid or C_6_H_8_O_6_ (Sigma, 33034), Perchloric acid or HClO_4_ (Carlo Erba, 409193).

HP-TLC: Toluene or C_6_H_5_CH_3_ (Sigma, 244511), Hexane or CH_3_(CH_2_)_4_CH_3_ (Sigma/Merck, 208752), Chloroform or CHCl_3_ (Sigma, 34854), Methanol or CH_3_OH (Sigma, 34860), 1-Propanol or CH_3_CH_2_CH_2_OH (Sigma, 402893), Ethyl acetate (VWR, 23880.290), Sulfuric acid or H₂SO₄ (VWR, 20700.298), Copper(II) sulfate pentahydrate or CuSO_4_·5H_2_O (Sigma, C8027), ortho-Phosphoric acid 99% or H_3_PO_4_ (Supelco, 1.00565.0500), magnesium chloride hexahydrate or MgCl_2_·6H_2_O (VWR, 25108.295). Lipid standards were acquired from Avanti® Polar Lipids – egg LPC, SM, egg PC, brain PS, liver PI, heart CL, egg PA, liver PE, egg PG, MSGCer (d18:1/17:0), GlucCer (d18:1/17:0), Egg Cer, MG 18:1, DG1.2 2xC18:0, DG1.3, TG 3×18:1, ChoE 17:0 – with the exception of Cholesterol (Sigma, C8667), ChoF (Merck, S448532) and AA (Sigma, A3555).

## High-performance thin-layer chromatography (HP-TLC)

Cells suspensions containing 8×10^6^ cells were divided into two equal parts to obtain cell pellets by centrifugation for either protein quantification or lipid extractions. Protein concentrations were determined as detailed above. For lipid extraction, cell pellets were extracted with the Bligh and Dyer method mentioned above with modifications. Quantification of phospholipids in total lipid extracts was based on the assay (Rouser et al., 1970)with minor modifications. An automatic TLC Sampler 4 (ATS4, CAMAG) was used to apply a total amount of 20 μg of phospholipid per sample in duplicates, lipid standards for identification and cholesterol standards (Avanti?) onto a Merck HPTLC glass plate silica gel 60 (20×10cm, layer thickness 200 µm).

HP-TLC was performed in sequential elution, the first 5 in an automated multiple development chamber (AMD2, CAMAG), then the following 3 in an automatic developing chamber (ADC2, CAMAG) with humidity control achieved with a MgCl_2_*5H_2_O solution. Solvent systems’ % distribution by order of elution: Ethyl acetate, 1-propanol, chloroform, methanol, 0.25% (W/V) aqueous potassium chloride, 1) 24:30:27:11:8, 2) 27:27:27:11:8, 3) 27:27:27:19:0, all up to 50 mm; 4) Ethyl acetate, chloroform (50:50), up to 55 mm; 5) Ethyl acetate, chloroform (30:70), up to 60 mm; 6) Hexane, ethyl acetate (60:40), up to 70 mm; 7) toluene up to 78 mm; 8) hexane up to 85 mm.

Plate surface was sprayed with a modified copper sulphate solution (Handloser et al., 2008) and revealed by heating at 100°C in a plate heater (CAMAG) for 25-30 min, then 5-10 min at 140°C. Imaging was done in a Fusion FX7 instrument (Vilber Loumat), using epi white light. Identification of the different classes of PL was carried out by comparison to PL standards applied in the TLC plate.

Plate images were analysed in Fiji, by determining peak areas corresponding to each lipid band of interest. Lipid distribution (%) of each lipid class was calculated per lane by dividing the peak area obtained for each band of interest by the sum of all peak areas of interest used in the same lane. Cholesterol quantification in each lane was calculated from a cholesterol standard quantification curve, then normalised to protein concentrations.

## Detailed lipidomics analyses

Acetonitrile (ACN) and isopropropanol (IPA) were purchased from Carlo Erba. Chloroform (CHCl_3_) was purchased from Merck. These solvents were LC-MS grade. Methanol (MeOH) was from VWR. Formic acid Optima LC-MS quality was purchased from Thermo Fisher Scientific. Ultrapure water was from Purelab flex (Veolia Water), and formate ammonium (99%) was from Acros Organics.

The extraction was performed using 1.5 mL solvent-resistant plastic Eppendorf tubes and 5 mL glass hemolyse tubes to avoid contamination. Methanol, chloroform and water should each be cooled down on wet ice before the lipid extraction.

A modified Bligh and Dyer was used for lipid extraction. One million cells were collected and pelleted in an eppendorf and 200 µL water was added. After vortexing (30s), the sample was transferred in a glass tube. 500 µl of methanol were used to rinse the Eppendorf tube and then transferred in the glass tube to which 250 µl of chloroform were finally added. The mixture was vortexed for 10 min before a new addition of 250 µl of water and 250 µl of chloroform. The mixture was vortexed again for 10 min and finally centrifuged (2500 rpm, 4°C, 10 minutes). Then, 400 µL of the organic phase was collected in a new glass tube and dried under a stream of nitrogen. The dried extract was resuspended in 60 µL of methanol/chloroform 1:1 (v/v) and transferred in an injection vial.

Reverse phase liquid chromatography was selected for separation with an UPLC system (Ultimate 3000, Thermo Fisher Scientific). Lipid extracts were separated on an Accucore C18 (150×2.1, 2.5µm) column (Thermo Fisher Scientific) operated at 400 µl/ minutes flow rate. The injection volume was 3 µL. Eluent solutions were ACN/H_2_O 50/50 (V/V) containing 10 mM ammonium formate and 0.1% formic acid (solvent A) and IPA/ACN/H_2_O 88/10/2 (V/V) containing 2 mM ammonium formate and 0.02% formic acid (solvent B). The step gradient of elution was in %B: 0.0 min, 35%; 0.0-4.0 min, 35 to 60%; 4.0-8.0 min, 60 to 70%; 8.0-16.0 min, 70 to 85%; 16.0-25.0 min, 85 to 97%; 25-25.1 min 97 to 100% B, 25.1-31 min 100% B and finally the column was reconditioned at 35% B for 4 min. The UPLC system was coupled with a Q-exactive Mass Spectrometer (Thermo Fisher Scientific); equipped with a heated electrospray ionization (HESI) probe. This spectrometer was controlled by Xcalibur software (version 4.1.31.9.) and operated in electrospray positive mode.

Data were acquired with dd-MS2 mode. MS spectra were acquired at a resolution of 70 000 (200 m/z) in a mass range of 250−1200 m/z with an AGC target 1e6 value of and a maximum injection time of 100ms. 15 most intense precursor ions were selected and isolated with a window of 1 m/z and fragmented by HCD (Higher energy C-Trap Dissociation) with normalized collision energy (NCE) of 25 and 30 eV. MS/MS spectra were acquired in the ion trap with an AGC target 1e5 value, the resolution was set at 35 000 at 200 m/z combined with an injection time of 80 ms.

Data were reprocessed using Lipid Search 4.2.21 (Thermo Fisher Scientific). The product search mode was used and the identification was based on the accurate mass of precursor ions and MS2 spectral pattern. The parameters for the ion identification and alignment were set up as follow: ion precursor tolerance 5 ppm; ion product tolerance 5 ppm; ion and adducts searched [H+], [NH4+] and [Na+]; alignment retention time tolerance 0.4 min; ID quality filters A, B and C.

## Supporting information

Source_file_2_lipidomics

Source_file_1_proteomics

## Expanded View Figures legends

**Figure EV1.**
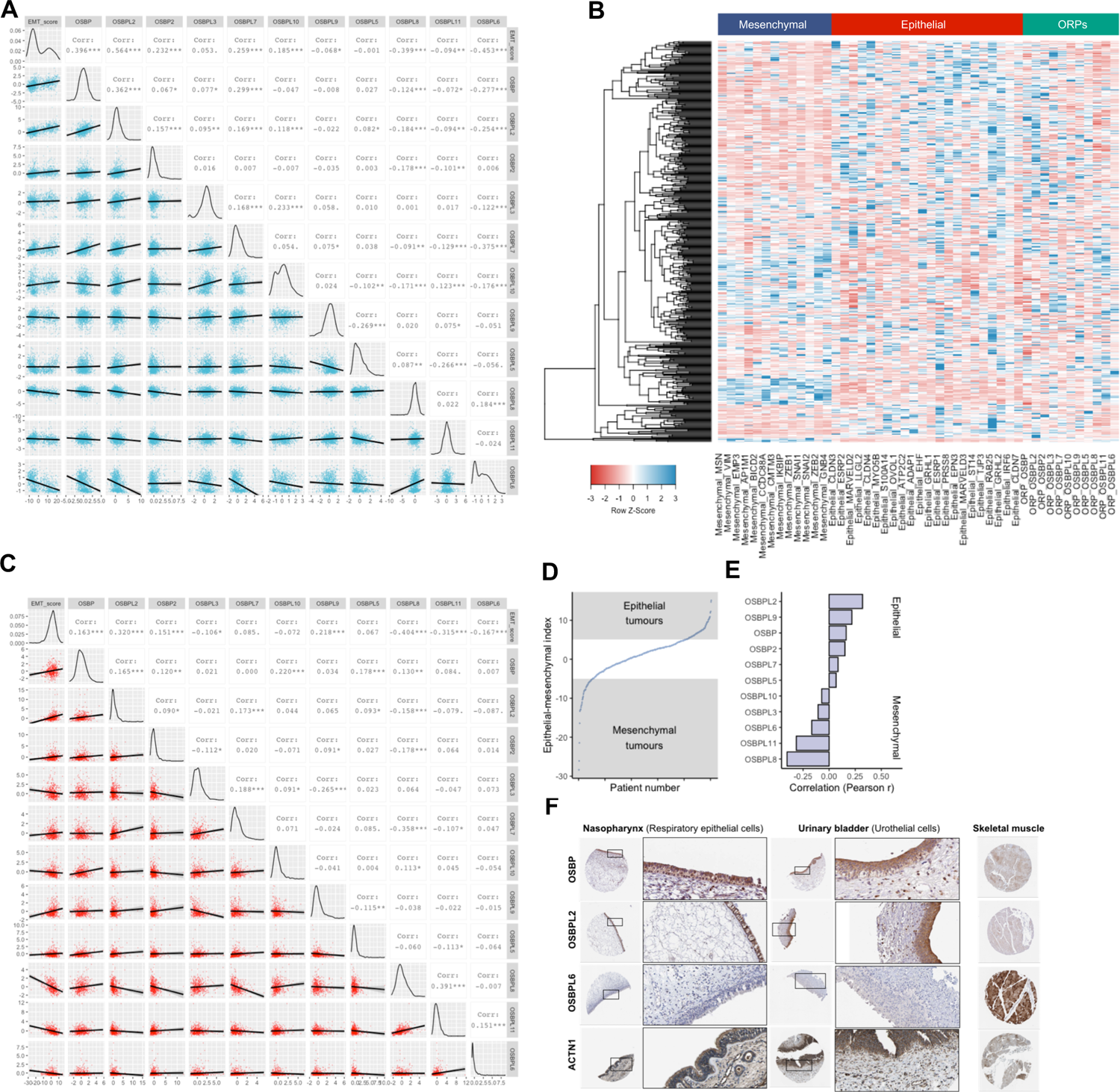
A. Correlation matrix comparing ORP expression values and epithelial-mesenchymal index corresponding to cell lines included in the Cancer Cell Line Encyclopedia. Corr=Pearson R, *P<0.05, **P<0.01, ***P<0.005.
B. Gene expression heatmap containing expression values for mesenchymal, epithelial marker genes as well as ORP genes of TCGA lung adenocarcinoma cohort.
C. Correlation matrix comparing ORP expression values and epithelial-mesenchymal indexes corresponding to tumour specimens included in the TCGA lung adenocarcinoma cohort. Corr=Pearson R, *P<0.05, **P<0.01, ***P<0.005.
D. Calculated epithelial-mesenchymal indices corresponding to each lung adenocarcinoma patient sample.
E. Correlations between epithelial-mesenchymal index and ORP expression values in TCGA lung adenocarcinoma samples.
F. Histology samples from The Human Protein Atlas show high epithelial-specific OSBP and OSBPL2 staining.

**Figure EV2.**
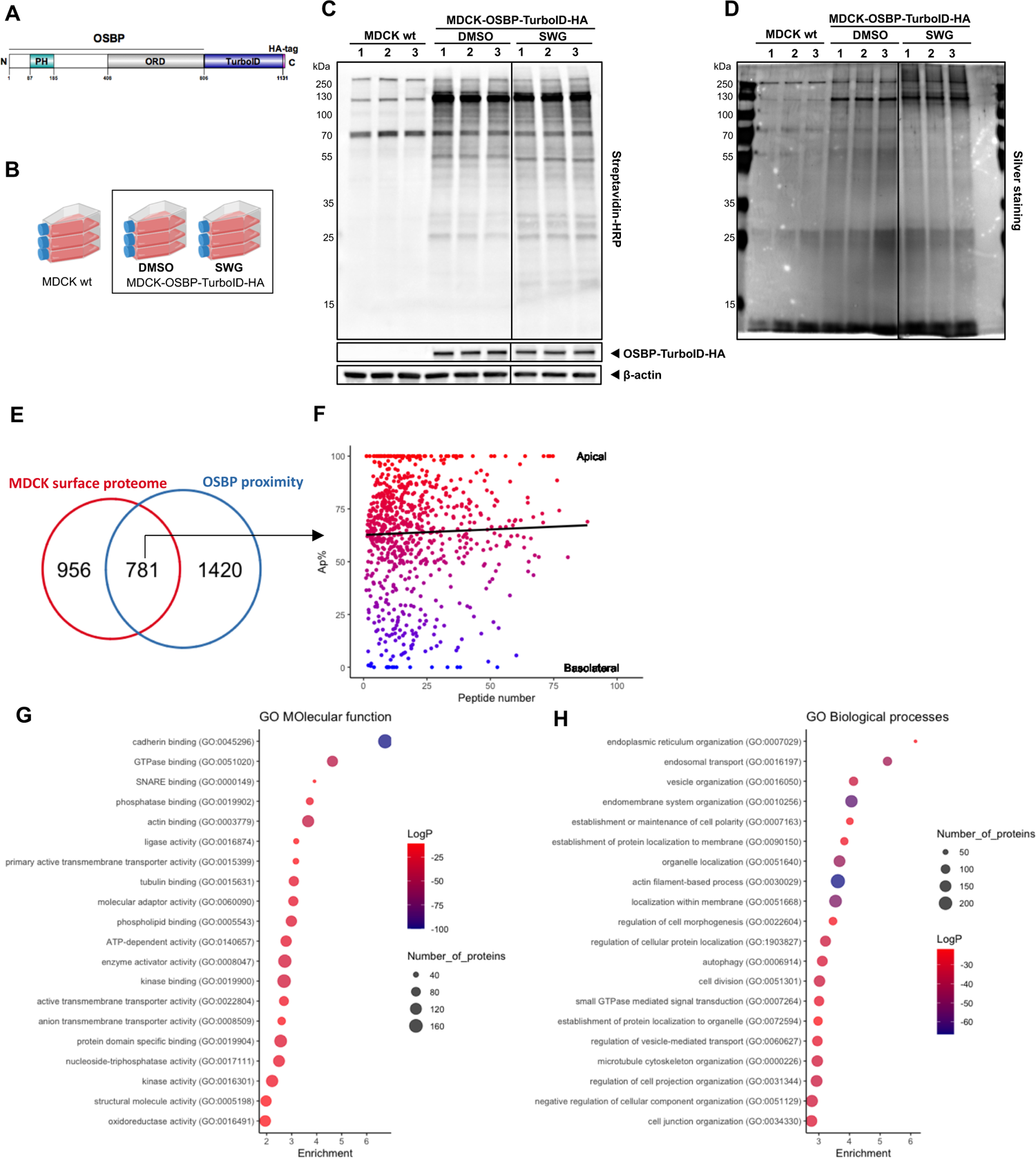
A. Domain structure of the OSBP-TurboID-HA construct stably expressed in MDCK cells.
B. Experimental design of the TurboID experiment.
C. Biotinylation profiles of cell lysates obtained from wild type MDCK or MDCK-OSBP-TurboID-HA cells. Membranes probed by streptavidin-HRP confirm successful protein biotinylation in MDCK cells stably expressing OSBP-TurboID-HA. Note that SWG-treatment affects the biotinylation profile.
D. Elution profiles of streptavidin beads incubated with lysates obtained from wild type MDCK or MDCK-OSBP-TurboID-HA cells. Biotinylated proteins were enriched on streptavidin beads, then beads were eluted by boiling, elution fractions were resolved on SDS-PAGE and revealed by silver staining. Elution profiles are comparable to biotinylation profiles shown on panel B.
E. 781 common hits were identified between the MDCK surface proteome (Caceres et al., 2019) and the OSBP proximity proteome (this study).
F. By analysing the overlapping hits, no correlation between apico-basal localization (Ap%) and the identified peptide numbers of the OSBP proximity partners can be established.
G. Enrichment analyses of GO terms classified by Molecular function.
H. Enrichment analyses of GO terms classified by Biological processes.

**Figure EV3.**
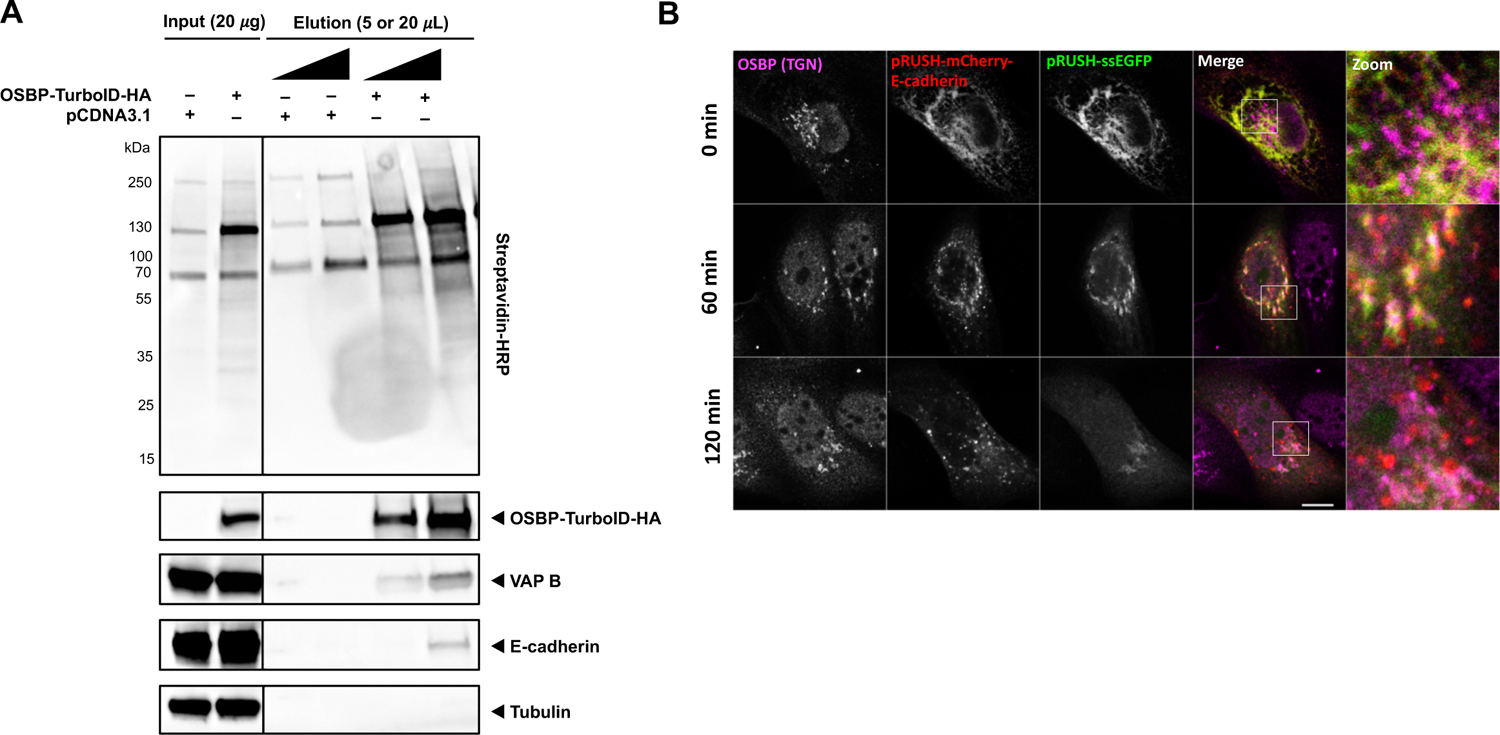
A. MDCK cells were transfected with control plasmid or with OSBP-TurboID-HA-expressing plasmid then biotinylated for 10 min on the next day. 5% of the cell lysates were kept for input control then biotinylated proteins were enriched on streptavidin beads. Beads were eluted by boiling, then samples were resolved on SDS-PAGE and transferred to membranes to probe with streptavidin-HRP or with specific antibodies. VAP B and Tubulin were used as positive and negative controls, respectively. Compared to the input samples, only a minor sample of the total E-cadherin pool was biotinylated.
B. Confocal microscopic analysis of MDCK cells co-expressing E-cadherin and ssEGFP RUSH cargoes and treated with biotin for the indicated times. Bar=12 #m.

**Figure EV4.**
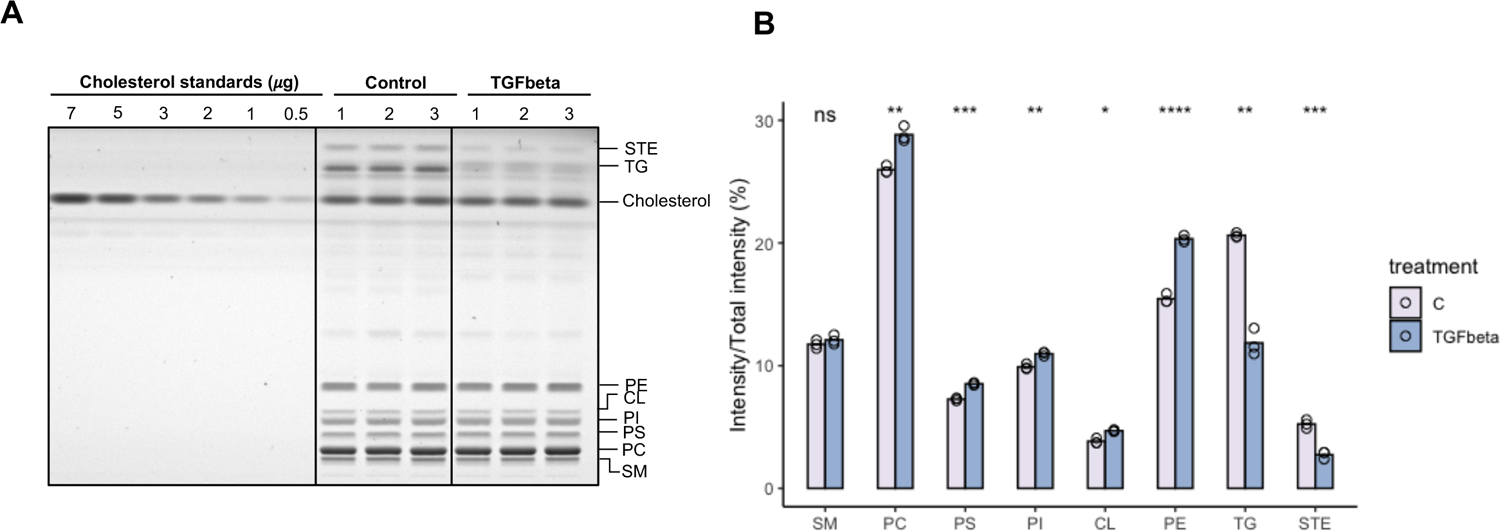
A. Photograph of the HP-TLC plate used for lipid class quantification. Abbreviation: STE – sterol esters, TG – triglycerides, PE – Phosphatidyl ethanolamines, CL – Cardiolipin, PI – Phosphatidyl inositols, PS – Phosphatidyl serines, PC – Phosphatidyl cholines, SM – Sphyngomyelines.
B. Quantification of lipid classes in control (C) and TGFbeta-treated A549 cells detected by HP-TLC. Bars indicate the mean of three independent experiments. *p<0.05; **p<0.01; ***p<0.005; ****p<0.001; ns=non-significant, Welch’s t test.

**Figure EV5.**
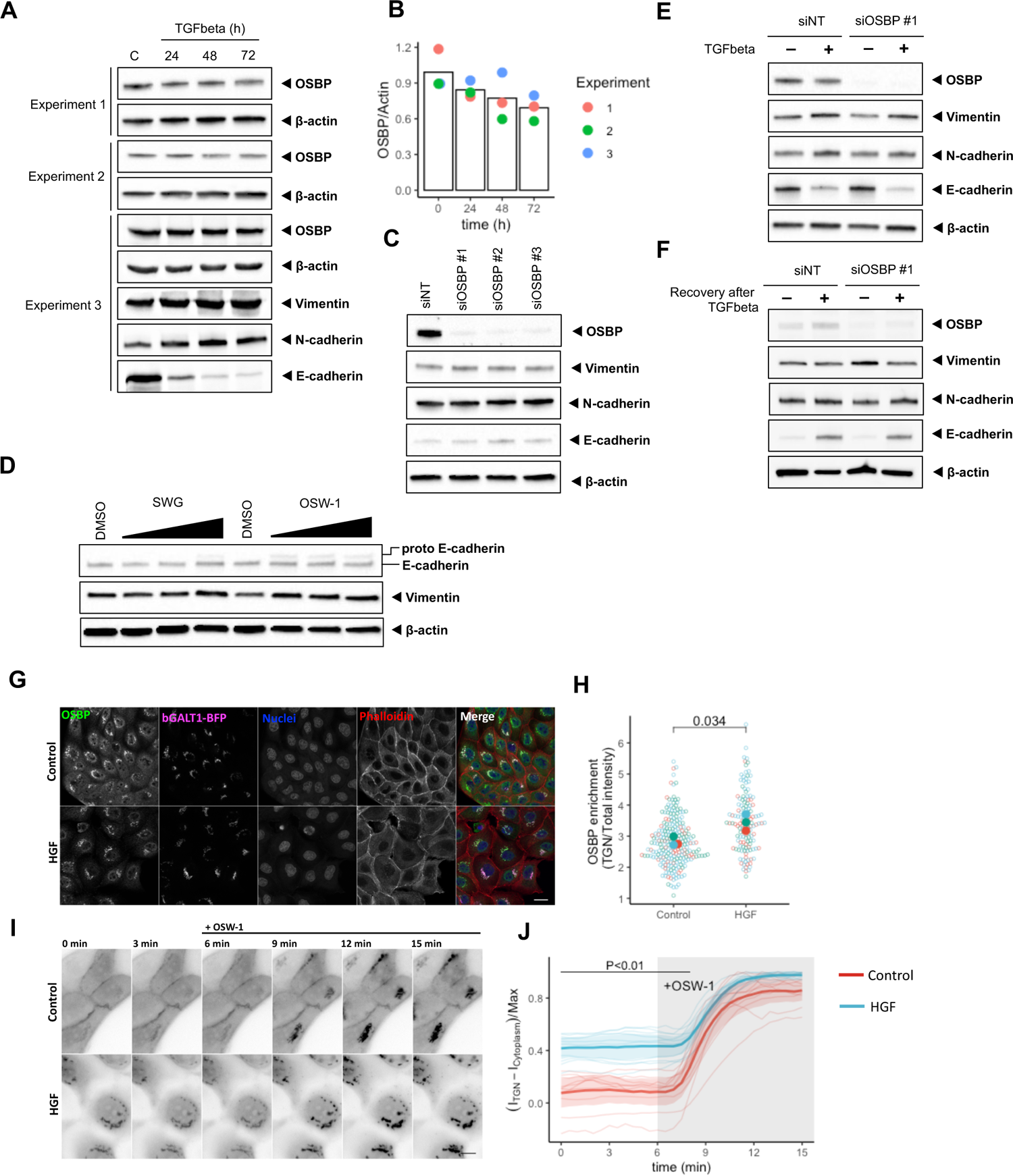
A. Western blot experiments showing the change of OSBP protein expression through EMT. 3 independent experiments are shown. E-cadherin was used as an epithelial marker, while N-cadherin and Vimentin served as mesenchymal markers.
B. Quantification of western blots shown in panel A.
C. siRNA-mediated silencing of OSBP does not affect EMT marker expressions.
D. A549 cells were treated 24 h with SWG and OSW-1 at 10; 100 an 1000 nM concentrations then E-cadherin and Vimentin levels were detected by western blotting. Uncleaved Pro-E-cadherin appears on the blots upon ORPphilin treatments.
E. TGFbeta can change the expression of EMT markers in the absence of OSBP.
F. A549 cells can re-express E-cadherin after washing-out TGFbeta-from the cells in the absence of OSBP as well.
G. OSBP is more enriched in the Golgi in HGF-stimulated MDCK cells compared to control cells. Bar= 20 #m
H. Superplot showing the redistribution of OSBP pools in HGF-treated MDCK cells. Means obtained from three independent experiments were compared by Welch’s t test.
I. (I-J) Kinetics of the GFP-P4M-SidM experiments in control and HGF-stimulated MDCK cells. One representative experiment is shown. OSW-1 was added to the cells in 50 nM concentration 6 minutes after starting the experiment. Each line corresponds to one cell and bold lines with shaded area indicate mean±95% confidence interval. Bar=10 #m

**Supplementary materials**

**Movie EV1**

**Movie EV2**

**Movie EV3**

**Movie EV4**

**Movie EV5**

**Movie EV6**

**Source Data 1 – Proteomics data**

**Source Data 2 – Lipidomics data**

**Supplementary Table**

## Acknowledgement

We are grateful for Frédéric Brau, Sophie Abélanet, Julie Cazareth and Nathalie Leroudier for their technical contribution. We thank Fanny Roussi for providing SWG. This work was supported by la Fondation pour la Recherche Médicale (DEQ20180339156) and the European Research Council (856404 — SPHERES). A post-doctoral fellowship to DK was provided by Agence National de la Recherche within the project entitled as Investissements d’Avenir UCA^JEDI^ (ANR-15-IDEX-01).

## Author contributions

DK, BA, BM and FL designed and conceptualised the research project. BA supervised the work. DK carried out the experiments. AP contributed to cloning design and molecular biology. ASG, LF and DD performed mass spectrometry, lipidomics and proteomics. ARDA performed lipid extractions and HPTLC. DK, BA analysed the data and wrote the manuscript.

